# Gut-associated functions are favored during microbiome assembly across *C. elegans* life

**DOI:** 10.1101/2023.03.25.534195

**Authors:** Johannes Zimmermann, Agnes Piecyk, Michael Sieber, Carola Petersen, Julia Johnke, Lucas Moitinho-Silva, Sven Künzel, Lena Bluhm, Arne Traulsen, Christoph Kaleta, Hinrich Schulenburg

## Abstract

The microbiome expresses a variety of functions that influence host biology. The range of functions depends on composition of the microbiome, which itself can change during the lifetime of the host as a consequence of neutral assembly processes, host-mediated selection, and/or environmental conditions. To date, the exact dynamics of microbiome assembly, the underlying determinants as well as the resulting effects on host-associated functions are not always well understood. Here, we used the nematode *Caenorhabditis elegans* and a defined community of fully sequenced, naturally associated bacteria to study microbiome dynamics and functions across the lifetime of individual hosts under controlled experimental conditions. By applying the neutral and null models, we demonstrate that bacterial community composition initially shows strongly declining levels of stochasticity, which, however, increase during late worm life, suggesting the action of random assembly processes in aged hosts following first colonization of *C. elegans*. The adult microbiome is enriched in strains of the genera *Ochrobactrum* and *Enterobacter* in comparison to the direct substrate and a host-free control environment. Using pathway analysis, metabolic, and ecological modelling, we further found that the lifetime assembly dynamics lead to an increase in gut-associated functions in the host-associated microbiome, possibly indicating that the initially colonizing bacteria are beneficial for the worm. Overall, our study introduces a framework for studying microbiome assembly dynamics based on the stochastic models and inference of functions, yielding new insights into the processes determining host-associated microbiome composition and function.

**Importance:** The microbiome plays a crucial role in host biology, with its functions depending on microbiome composition that can change during a host’s lifetime. To date, the dynamics of microbiome assembly and the resulting functions are not well understood. This study introduces a new approach to characterize the functional consequences of microbiome assembly by modelling both, the relevance of stochastic processes and metabolic characteristics of microbial community changes. The approach was applied to experimental time series data, obtained for the microbiome of the nematode *Caenorhabditis elegans*. The results revealed significant differences in host-associated and environmental microbiomes. Stochastic processes only played a minor role, and the host showed an increase in beneficial bacteria and an enrichment of gut-associated functions, possibly indicating that the host actively shapes composition of its microbiome. Overall, this study provides a framework for studying microbiome assembly dynamics and yields new insights into *C. elegans* microbiome functions.

## 1 Introduction

Microorganisms have shaped the evolution of multicellular host organisms since the very beginning [1]. As a consequence, they are of particular importance for the biology of their hosts. They can influence host nutrition and metabolism by providing specific enzymes for digestion (e.g., cellulase for digestion of plant material [2]) or certain nutritious compounds (e.g., specific amino acids [3]). They can define host development [4], maturation of the immune system [5], mediate immune-protection against pathogens [6], or even influence host behavior through interaction with the host’s nervous system [7]. During the lifetime of a host, the composition of its associated microbiome can change. For example, the assembly of the zebrafish gut microbiome is initially driven by neutral processes, while later time-points during development are characterized by non-neutral dynamics, most likely due to maturation of the immune system [8]. Overall, microbiome changes can be described by Vellend’s four high-level processes: selection, dispersal, drift, specification/diversification. The four processes help to generalize specific causes underlying microbial assembly, such as priority effects (positive-frequency selection), neutral assembly (drift, dispersal, specification), selection by the host, interaction of microbes with the host or with other microbes (selection) [9, 10]. To date, however, it is not well understood how these temporal changes affect the microbiome-mediated functions. If the host controls the microbiome-assembly process, then functional changes may be expected, because the host usually faces different functional challenges during its lifetime (e.g., development and immune protection early, sexual reproduction after maturation, etc). If the bacteria are in control, then later time points should be characterized by bacteria well adapted to the host habitat and/or highly competitive bacteria. A controlled experimental approach may help to better understand the consequences of microbiota assembly changes during host lifetime.

The nematode *Caenorhabditis elegans* provides a tractable experimental system, for which a large number of associated microorganisms have been isolated together with the nematode from the host’s natural habitat: rotting plant matter [11, 12]. Naturally associated microorganisms include members of the genera *Pseudomonas, Ochrobactrum, Enterobacter*, or *Comamonas*. Previous work highlighted that a representative set of the naturally associated microbes can provide almost all essential nutrients needed by the worm for growth (except cholesterol) and significantly affect *C. elegans* population growth [13]. Naturally associated bacteria can moreover provide immune protection against pathogens such as the Gram-negative *Bacillus thuringiensis*, the Gram-positive *Pseudomonas aeruginosa*, or the fungus *Drechmeria coniospora* [11, 14–16]. *Comamonas* gut bacteria influence dietary responses, development and life history of the nematode host [17, 18], while *Providencia* interacts with the *C. elegans* nervous system through production of the neuromodulator tyramine [19].

With the help of an experimental approach, microbiome compositional changes can be studied across time with the help of neutral assembly models and related to functions with the help of metabolic network models. In particular, we found surprisingly low neutrality for the microbial assembly in a former study which we intended to follow on longer time scales [20]. In addition, we recently developed a new method for metabolic pathway and model prediction [21] and proposed a classification of ecological strategies for microbes [13].

The aim of the current study is to use *C. elegans* and 43 bacteria representative of the nematode’s naturally associated microbiome (i.e., CeMbio43), in order to assess microbiome assembly dynamics and resulting functional consequences during the lifetime of the host. These 43 bacteria are an extension of the set of 12 bacteria that define the previously published CeMbio resource [22]. We obtained full genome sequences and reconstructed metabolic network models for these 43 bacteria. The bacteria were either combined with *C. elegans* or not (as control). Microbiome composition was characterized with the help of 16S rDNA amplicon sequencing for nematodes and the directly associated environment (labelled ‘substrate’) or the control environments without worms (labelled ‘control’) at 6 time points, covering *C. elegans* development and a considerable part of its adult lifetime (almost 8 days after egg hatching at 25 °C). We assessed diversity and neutrality of microbiome dynamics. We further used the reconstructed metabolic networks to infer changes in the metabolic competences of the microbial communities.

## 2 Material and methods

An extended version of material and methods can be found in the supplement.

### 2.1 Material

We used the *C. elegans* strain DH26, which is spermatogenesis defective and thus sterile at 25 °C, avoiding overlapping generations in our experiment, facilitating the repeated sampling across the lifetime of this nematode. We defined a community of 43 natural *C. elegans* microbiome isolates, the CeMbio43 (Table 1, Table S1, Table S2), which extends the previously described CeMbio resource (consiting of 12 bacterial strains [22]), which is representative of the natural *C. elegans* microbiome [11, 12], and which includes several strains for the naturally more abundant genera in the worm’s microbiome.

**Table 1:** Overview of the extended CeMbio community including genome statistics. Column ‘Ref’ indicates the reference studies which previously described the respective genomes: ‘*’ [22] and ‘**’ [13]. Genome size are given in Mega-base-pairs (Mb).

| ID | Species | Ref | Mb | Genes | Contigs | rRNA |
| --- | --- | --- | --- | --- | --- | --- |
| BIGb0170 | *Sphingobacterium multivorum* | * | 6.4 | 5391 | 1 | 21 |
| BIGb0172 | *Comamonas piscis* | * | 5.2 | 4595 | 1 | 19 |
| BIGb0393 | *Pantoea nemavictus* | * | 5.2 | 4667 | 2 | 22 |
| CEent1 | *Enterobacter cloacae* | * | 4.8 | 4458 | 1 | 26 |
| JUb134 | *Sphingomonas molluscorum* | * | 4.1 | 3816 | 4 | 9 |
| JUb19 | *Stenotrophomonas indicatrix* | * | 4.6 | 4079 | 2 | 14 |
| JUb44 | *Chryseobacterium scophthalmum* | * | 4.7 | 4199 | 1 | 21 |
| JUb66 | *Lelliottia amnigena* | * | 4.6 | 4207 | 1 | 22 |
| MSPm1 | *Pseudomonas berkeleyensis* | * | 5.7 | 5159 | 2 | 12 |
| MYb10 | *Acinetobacter guillouiae* | * | 4.6 | 4244 | 29 | 3 |
| MYb11 | *Pseudomonas lurida* | * | 6.1 | 5456 | 1 | 16 |
| MYb115 | *Pseudomonas fluorescence* |  | 6.3 | 5676 | 1 | 22 |
| MYb121 | *Erwinia billingiae* | ** | 5.0 | 4532 | 28 | 6 |
| MYb158 | *Acinetobacter johnsonii* |  | 3.6 | 3414 | 23 | 3 |
| MYb174 | *Enterobacter ludwigii* |  | 4.6 | 4293 | 29 | 5 |
| MYb176 | *Enterobacter ludwigii* |  | 4.6 | 4283 | 37 | 3 |
| MYb177 | *Acinetobacter sp. B2070* |  | 6.1 | 5449 | 50 | 4 |
| MYb181 | *Sphingobacterium faecium* | ** | 5.5 | 4619 | 243 | 8 |
| MYb186 | *Enterobacter sp. 638* |  | 4.8 | 4485 | 26 | 3 |
| MYb191 | *Acinetobacter sp. 4D-W-22* |  | 3.5 | 3350 | 21 | 11 |
| MYb21 | *Comamonas sp. B-9* |  | 5.4 | 4762 | 36 | 4 |
| MYb25 | *Chryseobacterium culicis* | ** | 5.0 | 4504 | 11 | 3 |
| MYb264 | *Chryseobacterium sp.* |  | 5.2 | 4607 | 1 | 21 |
| MYb328 | *Chryseobacterium sp.* |  | 5.6 | 5004 | 60 | 3 |
| MYb330 | *Pseudomonas sp.* |  | 6.1 | 5446 | 65 | 4 |
| MYb331 | *Pseudomonas sp.* |  | 6.1 | 5447 | 71 | 4 |
| MYb371 | *Pseudomonas sp.* |  | 6.1 | 5450 | 53 | 4 |
| MYb375 | *Erwinia sp.* |  | 4.9 | 4505 | 24 | 11 |
| MYb379 | *Ochrobactrum sp.* |  | 5.1 | 4780 | 47 | 5 |
| MYb382 | *Sphingobacterium sp.* |  | 4.7 | 4002 | 29 | 11 |
| MYb388 | *Sphingobacterium sp.* |  | 6.2 | 5326 | 29 | 3 |
| MYb396 | *Comamonas sp.* |  | 5.3 | 4719 | 23 | 3 |
| MYb398 | *Pseudomonas sp.* |  | 5.5 | 4895 | 105 | 4 |
| MYb416 | *Erwinia sp.* |  | 5.8 | 5380 | 443 | 14 |
| MYb49 | *Ochrobactrum anthropi* | ** | 4.8 | 4553 | 3 | 12 |
| MYb535 | *Erwinia sp.* |  | 4.9 | 4599 | 25 | 11 |
| MYb541 | *Pseudomonas sp.* |  | 6.3 | 5789 | 82 | 4 |
| MYb58 | *Ochrobactrum pituitosum* | ** | 5.0 | 4635 | 3 | 15 |
| MYb592 | *Gluconobacter wancherniae* |  | 3.2 | 3048 | 40 | 6 |
| MYb595 | *Gluconobacter cerinus* |  | 3.6 | 3299 | 48 | 2 |
| MYb596 | *Gluconobacter albidus* |  | 3.3 | 3044 | 51 | 3 |
| MYb69 | *Comamonas sp. TK41* |  | 5.3 | 4610 | 28 | 3 |
| MYb71 | *Ochrobactrum vermis* | * | 5.4 | 5191 | 3 | 12 |

### 2.2 Whole genome sequence analysis of bacteria

For the 26 bacterial strains without previously sequenced genomes (Table 1), we isolated high quality genomic DNA using a modified CTAB protocol, as described previously [13]. Short-read sequencing was performed by NextSeq 500 using the NextSeq 500/550 High Output Kit v2.5 (300 Cycles), and long-read sequencing with the Pacbio Sequel-54349U. The genomes were assembled using SPAdes v3.14.1 [23], MaSuRCA 3.4.2 [24], Unicycler v0.4.8 [25], shovill 1.1.0 [26], SKESA 2.4.0 [27]. Genome assembly quality was evaluated using QUAST v5.1.0rc1 [28]. The genome sequencing data was used to annotate genomic functions with gapseq 1.1 (Sequence DB: 1139b8e, bitscore cutoff 150; [21]), including inference of metabolic pathways, metabolic network models, growth media suitability, and carbon source and fermentation products. Virulence, resistance genes, and plasmids were scanned with abricate 1.0.1 [29] using the databases vfdb 2018-Jul-16 [30], plasmidfinder 2022-Mar-8 [31], resfinder 2022-Mar-8 [32], megares 2022-Mar-8 [33]. Gut microbiome specific gene clusters were identified by gutSMASH 1.0.0-1555cd7 [34]. For CAzyme annotation, dbCAN 3.0.2 was used [35]. The microbial interactions were simulated by a pairwise comparison of single vs. co-growth rate assuming TSB growth medium with BacArena 1.8.2 [36].

### 2.3 Analysis of microbiome changes

Temporal changes in microbiota composition were assessed for nematode populations (labelled ‘host’), the directly connected nematode growth medium (NGM) environment (labelled ‘substrate’), and the control NGM plates without worms (labelled ‘control’) in up to ten replicates and across six time points. DH26 *C. elegans* populations were synchronized the day before the experiment. Nematodes and bacterial lawns were sampled after 16 h (t2), 42 h (t3), 66 h (t4), 90 h (t5), 138 h (t6), and 186 h (t7). DNA was extracted with the Nucleo Spin® 96 Tissue Kit (Macherey-Nagel). The 16S libraries were prepared using the 341F and 806R primer covering the V3–V4 region of the 16S rRNA gene, followed by sequencing on the Miseq platform (Illumina). We processed the obtained 16S amplicon sequencing data using the standardized amplicon sequencing pipeline nf-core/ampliseq 2.1.1 [37]. Sample sequences were inferred by DADA2 [38], and the microbial composition data (ASV and taxonomic table) was analyzed with phyloseq 1.38.0 [39]. DESeq2 1.34.0 was used to find differentially abundant taxa [40] that were visualized by heat trees using the R package metacoder 0.3.5.001 [41]. To assess the neutrality of the samples the expected long-term distribution of a neutral model was fitted to the sample abundance data [20, 42]. To evaluate the impact of stochastic processes on community assembly, a null model called Normalized Stochasticity Ratio (NST) was evaluated using the R package NST 3.1.10 (function tNST; [43]). We combined the microbial 16S data with the functions predicted from genomic analysis and modeling. For each organism, the presence (true/false) of a particular function indicated if the organism’s abundance contributed to the abundance of the function. Finally, by adding up the abundances of all taxa in a sample for which the presence of a function was predicted, the overall abundance of the function was determined. In analogy to the taxon-level analysis, DESeq2 1.34.0 was then used to identify differentially abundant functions.

## 3 Data availability

The raw data from genome sequencing and 16S amplicon sequencing is available here [accession numbers link]. The source to reproduce figures and results can be found here [github link].

## 4 Results

### 4.1 The extended CeMbio43 community, genomes, and metabolic models

In 2020, we introduced a natural *Caenorhabditis elegans* microbiome resource, CeMbio, consisting of 12 representative members of the nematode’s native microbiome [22]. In the current study, we present an extended version of the CeMbio community with 43 bacterial strains, CeMbio43, which incorporates additional taxa chosen by their repeated presence in a long-term field study [12]. Newly integrated strains belong to the genera *Erwinia, Microbacterium*, and *Gluconobacter*. Besides a more comprehensive coverage of representative taxa of the native microbiome, the CeMbio43 community accounts for the redundancy detected for many microbiomes [44]. Additional strains to increase redundancy were added for *Acinetobacter* (+3), *Chryseobacterium* (+3), *Comamonas* (+3), *Enterobacter* (+3), *Ochrobactrum* (+3), *Sphingobacterium* (+3), and *Pseudomonas* (+6). Overall, the CeMbio43 community augments the existing resources for microbiome research in *C. elegans* by further improving the representation of a native microbiome and by accounting for species redundancy (Table 1, Table **??**, Table S2, Figure 1).

**Figure 1:**
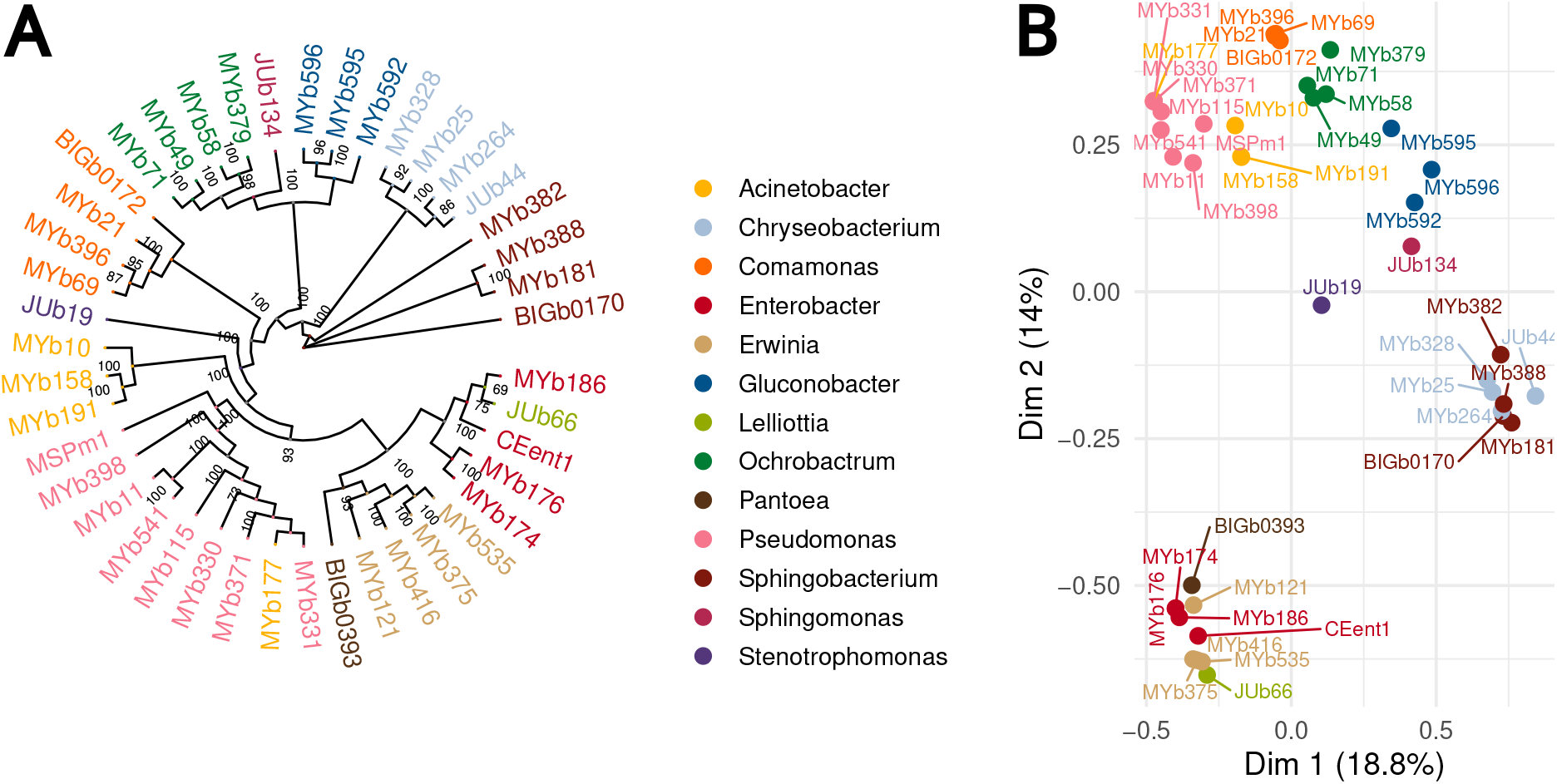
Genomes and inferred metabolic model similarities: (A) Rooted phylogenetic tree in circular format based on assembled genome similarity (alignment by gtdbtk and tree by iqtree, midpoint rooting), (B) Multiple Correspondence Analysis (MCA) of reactions that were present in the inferred metabolic models. The colors indicate the different genera of the CeMbio43 community. Strain codes are as in Table 1

For the 43 strains of the community, 12 genomes were assembled before for the original CeMbio community [22], and an additional five were already characterized for a previous study [13]. We now assembled the genomes for the remaining 26 strains. Noteworthy, depending on the organisms, the best assemblies were not obtained from one single but a combination of methods (MaSuRCA, shovill, SPAdes, Unicycler). SPAdes made more than 1/3 high-performing assemblies (Table S2). The CeMbio43 genomes were used to reconstruct a phylogenetic tree and, importantly, infer metabolic models and thus the metabolic competences of the included strains (Figure 1).

### 4.2 Microbial community changes across nematode lifetime

We studied changes in microbial communities from three environmental sources (control, substrate, host) for six time points (16, 42, 66, 90, 138, 186 h) with 8-13 replicates, totaling 180 samples. We measured the within-sample diversity and compared the influence of time and sample source on community dynamics, illustrated for the considered genera in the barplot of Figure 2A. The diversity increase over time was most robust for host and substrate samples (Spearman R=0.52; cf. R=0.32 (controls); Figure S7). The Shannon index showed significant temporal variation for substrate and host samples (Figure 2C). While a similar result was obtained for the evenness measure (Simpson), richness (Chao1) did not vary significantly over time (Figure S6). Analysis of the variance (ANOVA) of Shannon indices indicated the influence of sample source and time (Table S5). The Tukey Honest Significant Difference (Tukey HSD) Test suggested significant differences at the beginning and between 90 h and 186 h (Table S6) and for host vs. controls and host vs. substrate (Table S7).

**Figure 2:**
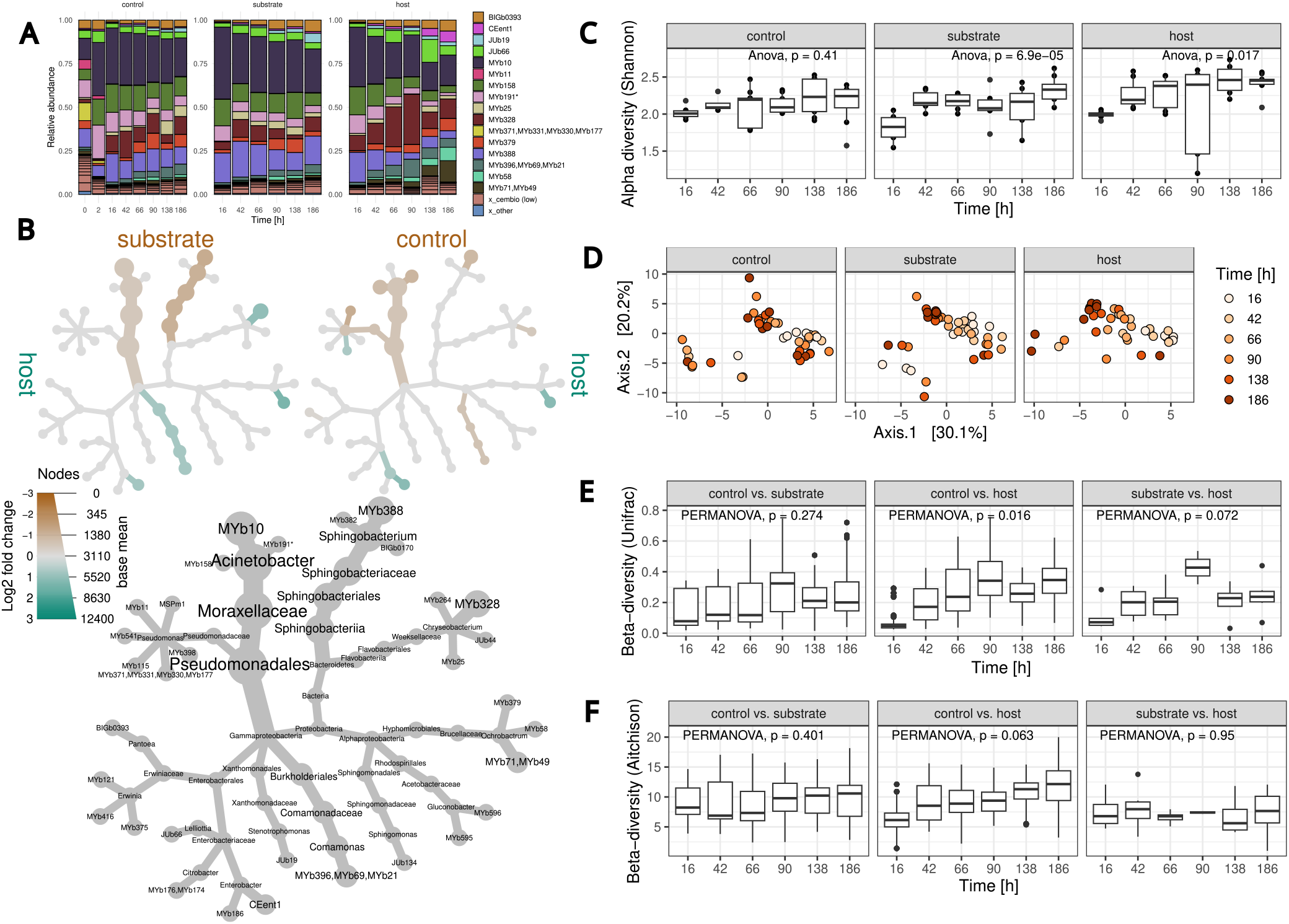
Microbial abundances and diversity across sample types and time. (A) Relative mean abundances per sample type and time point. Only samples with a maximal relative abundance *>*= 0.05 are shown. (B) Differential heat tree of species abundances. A taxonomic tree is shown in gray. The colored trees compare conditions: substrate vs. host (top left), control vs. host (top right). Differentially abundant species were colored brown (enriched in substrate or controls) or turquoise (enriched in host). Differential abundances were inferred by DESeq2. Taxa with adjusted P-value of *≤* 0.05 are shown. The thickness of branches corresponds to mean abundance. (C) *α*-diversity of filtered and agglomerated microbial samples on taxon level (Shannon diversity) across time. (D) Ordination of microbial samples shown in a PCoA plot using Aitchison distance. Time is indicated by increasing red color intensities. (E),(F) Differences in *β*-diversity (Aitchison, weighted unifract distance) compared between sample types and across time. For comparing host-associated samples, pairs from the same replicate were available, whereas for comparing host with control or substrate with control, pairs were randomly associated (100 repetitions).

When comparing differences between samples (*β*-diversity), we found the influence of time on sample variation to be most prominent for host-derived samples (Aitchison distance; Figure 2D). Although we found significant differences between sample sources (PERMANOVA F test P=0.002, Table S8), variation over time was only identified for host and substrate samples (PERMANOVA F test P*<* 0.05 for Bray-Curtis, Unifrac, and Aitchison distance; Figure S8, Table S10). Time points 66 h and 186 h showed significant dissimilarities across sample sources (Table S9). For some samples, pairs from the same experiment existed so that sample distances could be compared directly. We observed that differences between host samples and controls increased over time (Figure 2F, S9). For host and substrate samples, Bray-Curtis and Unifrac distances indicated that initially diverge in composition, but after 90 h, become more similar again (Figure S9, 2E).

We performed a differential abundance analysis using DEseq2 based on a negative binomial generalized linear model with time and source as variables to identify bacterial strains whose growth showed variation. The results were summarized in a ‘heat tree’ that depicts differentially abundant taxa for host vs. control and host vs. substrate. When comparing control vs. substrate samples, we did not find differentially abundant taxa (Figure 2B). In agreement with previous studies, *Ochrobactrum* (MYb71/MYb49) and *Enterobacter* (CEent1) strains were exclusively enriched in samples obtained from within the host in all comparisons. *Chyseobacterium* (MYb328) and *Pseudomonas* (MYb371/MYb331/MYb330/MYb177) showed higher abundances in host compared to control samples, whereas *Comamonas* (MYb396/MYb69/MYb21) and *Chryseobacterium* (MYb328) were more abundant in host compared to substrate samples. *Acinetobacter* (MYb10, MYb158, MYb191) belonged to the most widely abundant taxon across all samples and was, together with *Sphingobacterium* (MYb388), significantly enriched in control and substrate samples. In addition, *Pseudomonas* (MSPm1), *Chryseobacterium* (JUb44), and *Sphingomonas* (JUb134) showed higher abundances in controls in contrast to host samples (Figure 2B, Figure S5).

In summary, we found differences in diversity between samples obtained from hosts, substrate, and controls. The effect of time was most pronounced for microbial samples obtained from the host, and host samples diverged over time compared to controls. This suggests that specific microbial taxa were enriched in the host-associated community compared to those from substrates or controls.

### 4.3 Stochasticity

We next assessed to what extent deterministic and stochastic processes have shaped the assembly of microbial communities. We first took Hubbell’s neutral model to investigate the effect of stochastic effects. In a previous study, we observed low neutrality for the microbial community assembly of *C. elegans* compared to other organisms (goodness of fit *R*^*2*^ = 0.44). Since the fit of the neutral model is based on the long-term equilibrium state, we hypothesized that the microbial community in the short-lived nematode might still be in a transient state on its way to a more neutral composition, thereby underestimating neutral effects [20]. Therefore, in the current study, we characterized the microbial community dynamics over *C. elegans* lifetime and tested whether neutrality increases across time. Surprisingly, we found that the neutral expectation fitted the observed community abundance data very well (0.8 *< R*^*2*^ *<* 0.95) regardless of the sample source, indicating highly neutral dynamics (Figure S11). Although we did not find a continuous increase in neutrality, the dispersal parameter m of the neutral model decreased over time and differed between sample sources (Figure 3A). Lower values for the dispersal parameter at later times could indicate less mixing between samples, leading to more variance between communities. However, the neutral model analysis comes with the limitation that it is based on the presence and absence of taxa, while most samples of our study continuously contained nearly all of the CeMbio43 taxa and only their relative abundance varied. Because of this limitation, we further assessed the role of stochastic effects with the taxonomic normalized stochasticity index (tNST). The tNST quantifies the effect of deterministic (tNST*<* 0.5) and stochastic effects (tNST*>* 0.5) based on beta-diversity. The tNST changed over time for all conditions, starting with higher values (tNST*>* 0.5) followed by a decrease, most prominently for host samples (tNST*<* 0.2 after 90 h), and a late increase observed only for host samples (*tNST ≈* 0.4 after 186 h; Figure 3C). In general, host samples showed significantly lower stochasticity values when compared to susbstrates and controls (Figure 3B). Taken together, contrary to our original hypothesis, stochasticity rather decreases across time, even if there is a slight reversal at the late time points for the host samples. This may suggest that non-stochastic processes shape composition of the microbial communities in our experiment.

**Figure 3:**
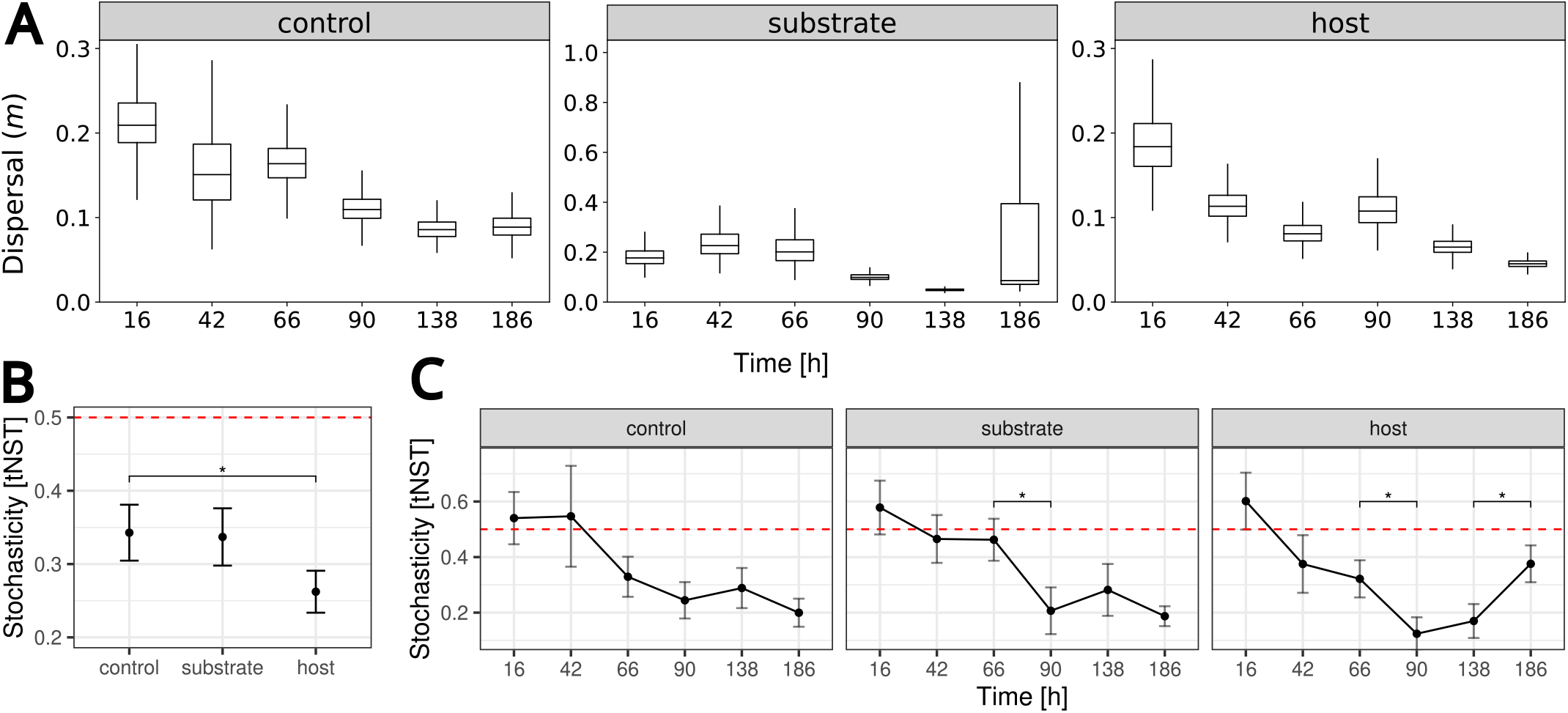
Influence of stochastic effects on microbial community dynamics. (A) Best-fit values of the dispersal parameter *m* of the neutral model across time and the different sample types. (B),(C) taxonomic Normalized Stochasticity index (tNST) across time and sample types. Red dashed line (tNST= 0.5) indicates the transition of the influence of stochastic (tNST*>* 0.5) to deterministic (tNST*<* 0.5) processes in community assembly. Significant differences in tNST was observed between sample types (panel B) and across specific adjacent time points (panel C)

### 4.4 Enrichment of host-specific functions

Our diversity analysis and the examination of stochasticity indicated the presence of deterministic effects on microbiome assembly and differences in these dynamics between sample types (control, substrate, worm). Therefore, we wondered whether the non-stochastic effects are explained by the selection of specific metabolic functions of the microbiome across time. To address this idea, we assessed the temporal changes in metabolic characteristics of the microbial community, as inferred from the microbiome abundance data in combination with the corresponding genome sequences and recontructed metabolic models. For our analysis, we considered distinct types of microbiome characteristics (see Methods), including those defined by carbohydrate-active enzymes (i.e., cazyme in Figure 4), metabolic exchange (i.e., exchange), microbiome-related gut functions (i.e., gut), pair-wise interactions (i.e., interactions), growth media (i.e., medium), metabolic pathways (i.e., metabolism), adaptive strategies (i.e., uast), and virulence factors (virulence). The predicted characteristics (presence/absence) were related to the microbial abundance data to obtain temporal information for each potential function and community.

**Figure 4:**
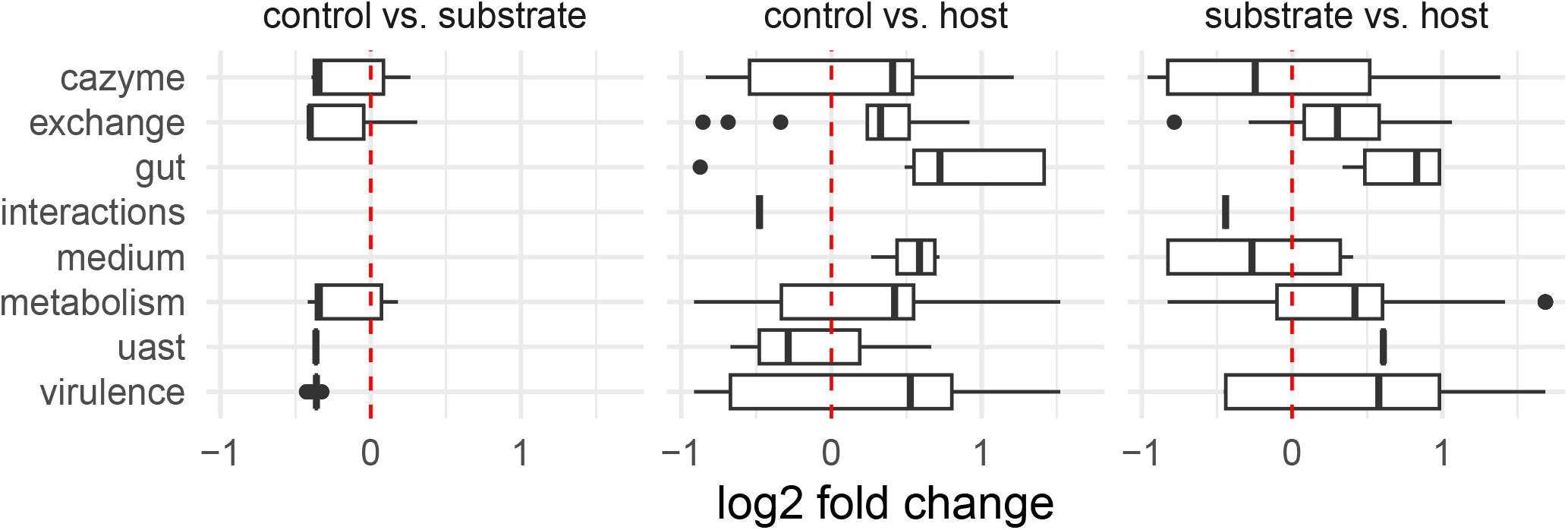
Differentially abundant functions in either host, substrate, or control microbiome. Differential abundance was assessed by DESeq2 for different functional subsystems (along vertical axis) and different comparisons of the sample types (the three panels). For example, for the comparison ‘substrate vs. host’, positive values (log2 fold change) indicate that the host-associated microbiome showed an increased presence of the functions of the respective subsystem, while negative values a comparative decrease of the functional subsystem in the host samples. The following subsystems were considered: Cazyme: carboydrate-active enzymes. exchange: carbon/energy source and metabolic byproducts. Interactions: prevalence of ecological interactions based on pairwise metabolic predictions. Medium: growth medium components. Metabolism: MetaCyc pathways. Microbiome: gut-microbiome gene clusters. Virulence: Virulence and resistance genes.

Our analysis revealed that the functional diversity was highest and increased over time for worm-associated microbial communities (Figure S13). We then performed a differential abundance analysis using DESeq2 and observed substantial trait differences for many comparisons between sample types (Figure 4). We next wondered which exact metabolic functions underlie the observed differences for individual microbiome characteristics. For carbohydrate-active enzymes, we used the automated CAzyme annotation dbCAN pipeline to predict carbohydrate-active enzymes and found glycoside hydrolases and glycosyl transferases to be differentially abundant between sample sources (Figure S14). The corresponding substrates mainly comprised oligosaccharides and o-glycosyl compounds (Figure S15). As a next step, we inferred substrate uptake and metabolic products from metabolic models (Figure S16). Among others, we found the capacity for tryptophan uptake and folate production to be enriched in the host-assocaited microbiota. Furthermore, gene clusters responsible for gut microbiota functions (predicted by gutSMASH) were almost exclusively enriched in host samples. This result was due to an increased abundance of genes encoding for short-chain fatty acid production, polyamine metabolism, and anaerobic energy metabolism (Figure 5).

**Figure 5:**
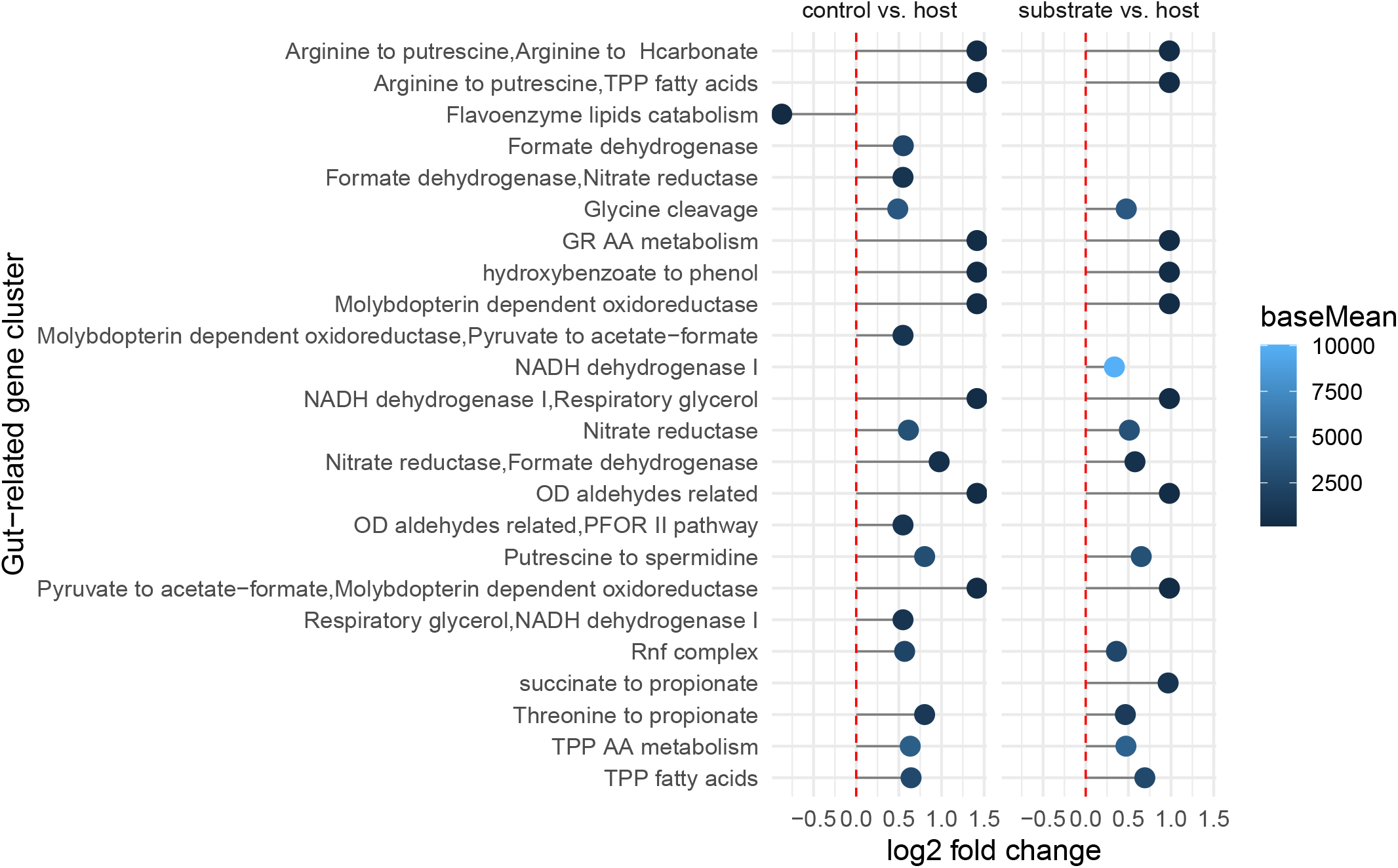
Enrichment of gut microbiome functions. Gene clusters were predicted by gutSMASH and combined with microbial abundance data. We almost exclusively found gut functions enriched in host samples by differential abundance analysis. No differences were found for controls vs. substrate. Abbreviations: thiamine pyrophosphate (TPP), amino acids (AA), glycyl radical amino acid metabolism (GR), oxidative decarboxylation (OD), pyruvate ferredoxin oxidoreductase (PFOR)

We also used the metabolic models to infer pair-wise interactions between bacteria. We found neutral interactions to be enriched in non-host samples (Figure S17). In addition, we predicted the suitability of different growth media or nutrients from metabolic models. We found that host-enriched microbes could more often use sulfoquinovose and, to a lesser extent, N-acetylglucosamine and N-acetyl-neuraminate (Figure S18). We further studied the presence of metabolic pathways as defined by MetaCyc (Figure S19). An over-representation analysis identified pathways involved in secondary metabolites degradation (host vs. substrate) and cofactor biosynthesis (host vs. control) enriched for host samples. Samples from substrates showed fatty acid and lipid degradation (substrate vs. worm), and control samples had increased aromatic compounds and amine degradation (control vs. worm) (Table S11).

We next assessed differential abundance of specific pathways known to be relevant for *C. elegans*. Here, vitamin B12 biosynthesis and salvage pathways were differentially more abundant in the host than in substrate samples (Figure S23). Moreover, branched-chain amino acids degradation pathways were enriched in the host compared to control samples (Figure S24). In a previous study [13], we showed that it could be informative to classify bacteria into competitors, stress-tolerator, and ruderals, according to the universal adaptive strategies theory (UAST) [45]. Among traits relevant to adaptive strategies, we found an enrichment of codon-usage bias in host samples, biofilm production (extracellular polymeric substances, eps) in control samples, and in general, less competitive traits in substrate samples (Figure S20). We also explored the differential abundance of genes defined by the virulence factor database. We identified motility and biofilm genes enriched in controls and immune regulation enriched in host samples (Figure S21). Last, we checked explicitly for two pathways relevant to host colonization and worm fitness as identified in our previous study [13]. Consistent with our previous results, we found pyruvate fermentation to acetoin and hydroxyproline degradation almost exclusively to be differentially abundant in the host-associated microbiota (Figure S22).

As a final step in our analysis, we tracked the contribution of microbial species to the inferred functions. For this, we summed up the species abundances and performed a regression analysis to account for the impact of low-abundant species on functional abundances. Although some taxa showed a more substantial contribution to some functions (as for example seen for the *Enterobacteriaceae* CEent1 and JUb66), we found that the enrichment of gut functions was supported by a larger variety of microbiota strains (Figure 6, S25).

**Figure 6:**
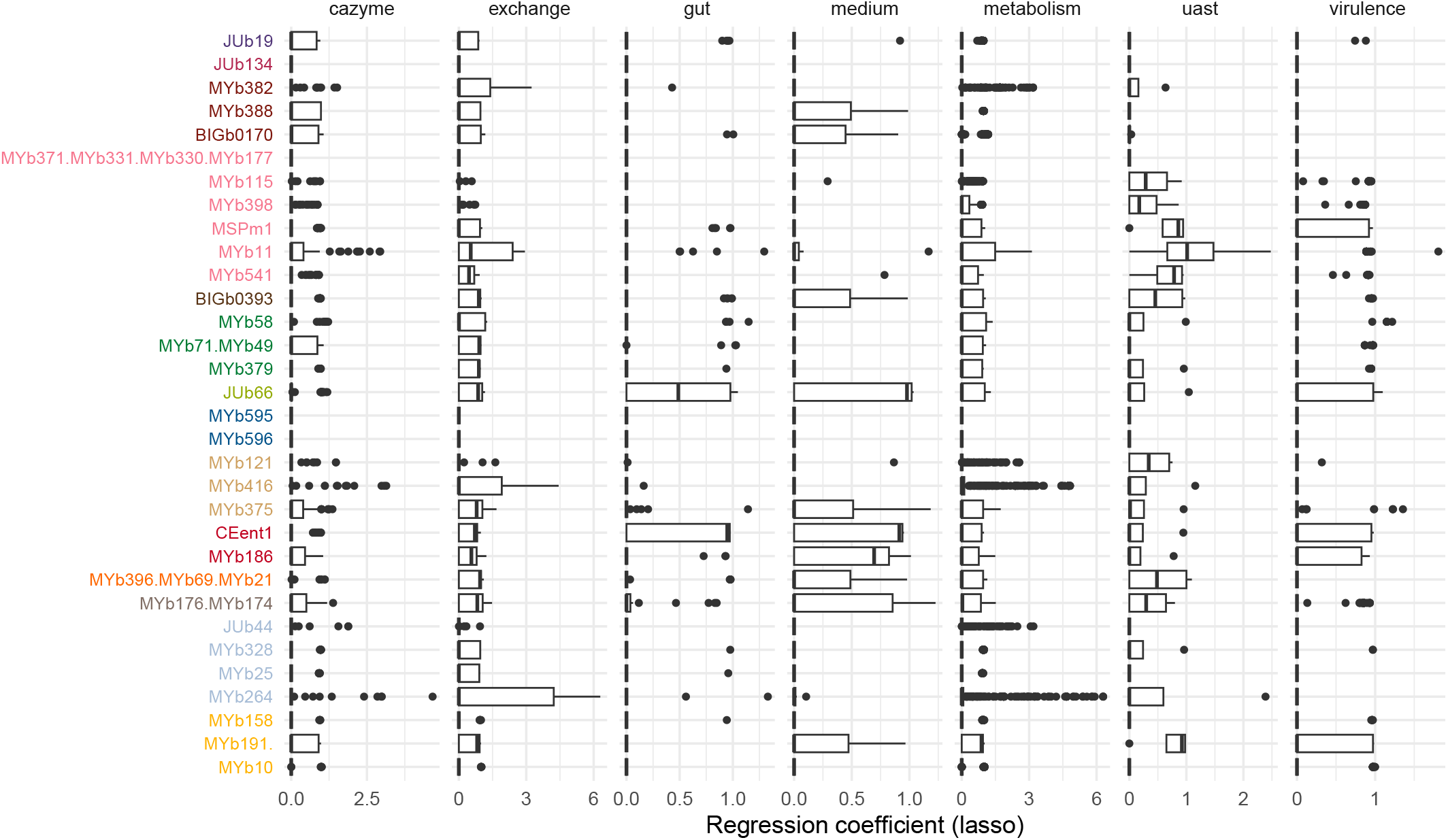
Contribution of microbial taxa to functions. The attribution of individual taxa to predicted functions were determined by regression analysis. We used a penalized regression model to estimate how well the taxon abundances can predict functional abundances. For each taxon, all coefficients from regression models summarized by subsystem are shown in boxplots. The subsystem ‘interactions’ is not shown because of low variance. Strain codes are as in Table 1; and colors indicate different genera as in Figure 1

Taken together, we identified many potential functional traits that differ between sample types. The microbial communities from *C. elegans* were signficantly enriched for traits relevant to a host environment.

## 5 Discussion

In our study, we analyzed the assembly of a newly defined and fully sequenced microbial community across a substantial part of the lifetime of *C. elegans*. Our study is one of very few studies to characterize microbial community dynamics in this nematode across time. We compared microbiome characteristics across seven time points and a total of 186 hours for three different sample types: host, the direct substrate environment of the hosts (substrate), and an environment without nematode hosts (controls). We identified that time affected alpha diversity only for substrate and host samples. The microbial composition of host vs. control samples became less similar over time, indicating the relevance of host presence. We further identified taxa which were specifically enriched in host samples, namely strains from the genera *Ochrobactrum* and *Enterobacter*. We also found comparatively little influence of stochastic effects on microbial assembly across our sample types, including the host samples, possibly suggesting deterministic processes shaping microbiome composition. Finally, we inferred changes in microbial functions from the fully sequenced genomes and community composition and identified several functions, which were previously described for host-microbe interactions, to be specifically enriched in the host-associated microbial communities.

### 5.1 Specific beneficial microbes increase in abundance in the host

In our previous study on the 12-member CeMbio community, we found *Ochrobactrum* (MYb71) and *Stenotrophamonas* (JUb19) more abundant in worms, while *Acinetobacter* (MYb10) and *Sphingobacterium* (BIGb0170) were enriched in lawns [22]. Consistent with these previous results, we here confirm a higher abundance of *Ochrobactrum* (MYb49/MYb71) in worms, and of *Acinetobacter* (MYb10, MYb191, MYb158) and *Sphingobacterium* (MYb388) in the host-free controls. In contrast to the earlier results, we now also found *Enterobacter* (CEent1) to be more abundant in worms, but not *Stenotrophamonas*. The latter differences may be explained by the more diverse CeMbio43 community, the different *C. elegans* strain used, and/or the higher temperature of our experiment (which was necessary to ensure worm sterility and thus unconstrained sampling of worms across its lifetime). Especially if other bacterial species of the same genus, as in the case of *Sphingobacterium*, followed the same enrichment pattern, the extended CeMbio community’s additional redundancy might help to identify the species or strains performing the same role or function. For the host-enriched *Ochrobactrum*, beneficial effects for *C. elegans*, such as ROS detoxification and increased growth, were reported before [46–48]. In addition, *Enterobacter* bacteria were shown to have an immune-protective effect in *C. elegans* [49]. The combination of known positive effects and the increased abundance of *Ochrobactrum* and *Enterobacter* in the nematode gut may suggest that *C. elegans* is able to favor presence of beneficial microbes. A dissection of the possible underlying molecular mechanisms clearly warrants further research.

It is possible that the nematode immune system can shape the composition of the associated microbial community. Indeed, a previous comparison of microbiome composition across a diverse set of natural *C. elegans* strains identified distinct microbiome types, most likely as a consequence of host filtering, and demonstrated a significant role of worm genetics in this context [48]. Interestingly, in this previous study, host filtering favored presence of *Ochrobactrum* and this effect involved the worm’s insulin-like signaling pathway [48]. Although we did not find a specific *Ochrobactrum*-dominated microbiome type, *Ochrobactrum* was still significantly more abundant in worms compared to controls or substrate samples (Figure S5). Moreover, our analysis of microbiome-associated functions yielded a possible link to host insulin signaling. In detail, the host samples with their high *Ochrobactrum* abundance were significantly enriched for the microbiome-associated branched-chain amino acid (BCAA) degradation pathways (Figure S24), which was previously reported to contribute to insulin resistance [50], and a reduced catabolism of BCAA increased the lifespan of *C. elegans* [51].

Our results may further shed light on the characteristics of a possible core microbiome of *C. elegans*, as previously proposed by colleagues and us with the help of a meta-study [52], even though more recent field data suggests high variability in the presence of putative core taxa across time [12]. In general, the precise definition and quantification of a core microbiome remain challenging [53]. Therefore, Turnbaugh and colleagues argued that a core microbiome exists primarily on the level of shared metabolic functions [44]. In our study, we introduced a microbiome community that is representative of the microbial diversity encountered in *C. elegans* in nature and that therefore is already close to what may be considered a core microbiome of this nematode. We still observed variation in microbiome composition across time and sample type. Importantly, we found the presence of host-relevant functions to be enriched in the host samples and distributed across numerous microbial taxa (Figure 6). This finding is consistent with the proposed importance of functions rather than specific taxa in defining the host-associated core microbiome [54].

### 5.2 *C. elegans* microbiome assembly dynamics are shaped by selective rather than stochastic processes

Stochastic processes have been reported to shape the assembly of microbial communities in diverse contexts and hosts systems [55]. Previously, we used the neutral model to assess stochasticity in the microbiomes of several different hosts and found the lowest neutrality for the microbiome of *C. elegans* [20]. Considering the short lifespan of the nematode, we previously hypothesized that the worm-associated community might be in a transient state that can appear non-neutral temporally. In the current study, we now observed a much higher level of neutrality, even though we considered a longer time-span, with very little change in neutrality over time.

However, the *α*-diversity of samples showed differences only for evenness rather than richness (Figure 2C, S6), indicating little variation in the general presence of taxa. This makes sense and may have even been expected since we used a pre-selected community of microbes known to coexist with *C. elegans* in nature. Consequently, the classical neutral approach was limited because it is based on differences in presence-absence as a function of mean relative abundances, while our data set shows little variation in presence/absence of the bacteria, but instead is rather shaped by changes in relative abundances.

As an alternative to the neutral model, we therefore employed a null model approach, which is less influenced by presence/absence of microbes and has shown higher accuracy and precision than the neutral model in certain cases [43]. The null model analysis revealed a decrease in stochasticity for all environments initially, followed by increased stochasticity for later time points only for host samples. Initially, the observed stochasticity dynamics could be a consequence of the experimental setup for which abundances were arranged artificially and did not reflect ecological conditions. Therefore, an initial decrease in stochastic effects seems reasonable to establish the more natural strain abundances and may be further influenced by selective processes in the host. The later increase of stochasticity only for host samples is generally consistent with previous reports from other animal hosts [20] and most likely reflects the aging of *C. elegans*, in which the constant pumping of new material and grinder efficiency declines after around 100 hours [56]. Consequently, the host is likely to lose its ability for selecting the presence of microbes, so that microbiome composition is more strongly influenced by stochastic processes. Such an age effect has also been reported for the human microbiome, where the microbial community composition becomes less diverse and more unique with aging [57, 58]. Although stochasticity was highest for host samples at later time points, the overall stochasticity (i.e., summarized across time) of host-derived samples was significantly lower than that of control and substrate samples. Given the similar starting point of stochasticity for all environmental sources, we conclude that deterministic (i.e., selective) processes may play a particular role in shaping host-associated microbial communities, consistent with the previous demonstration of host genetic influence on microbiome composition in *C. elegans* [48].

### 5.3 Host-relevant traits become enriched in the *C. elegans* microbiome

We used our time-series abundance data in combination with the genome sequence-inferred metabolic characteristics to determine which microbial traits predominate in the different sample types. For the host samples, we found significantly higher abundance of many gene clusters and pathways that are known to be produced by gut microbes and affect animal hosts [59]. One example is the enriched production of short-chain fatty acids, which can have many beneficial effects in animals, including protection of *C. elegans* against neurodegeneration [60, 61]. High levels of the microbial derived short-chain fatty acid propionate can also have a toxic effect in *C. elegans* and its degradation is linked to vitamin B12 whereby vitamin deficiency reroutes the fatty acid metabolism [18, 62]. Our former study identified several members of the native microbiome of *C. elegans* to possess pathways for vitamin B12 biosynthesis [13]. In agreement with these findings, we found propionate and vitamin B12 production capacity enriched in microbial samples from *C. elegans*, indicating a vitamin-rich environment with B12-dependent host breakdown of propionate [62].

Another example is the higher production of polyamine spermidine and folate in the host-associated microbiome. Spermidine is known to extend the lifespan of *C. elegans* and folate is an essential vitamin for this nematode, which affects germ cell proliferation and ageing [63, 64] and thus fitness-associated characteristics. In addition, the host-derived microbial samples showed higher potential to degrade N-acetylglucosamine and N-acetyl-neuraminate used for the dense glycosidation of host mucins and thus provide a source for host selection of commensal bacteria [65]. However, *C. elegans’* glycans seem to lack N-acetylneuraminate [66]. Nonetheless, *C. elegans* contains gangliosides [67] that could provide N-acetylglucosamine because gangliosides are glycosphingolipids that contain N-acetylglucosamine as terminal residue [68]. Furthermore, we looked into life-history traits and found a codon usage bias for bacteria enriched in *C. elegans*. A codon usage bias is a form of genomic adaption that changes gene expression, thus permitting microbes to adapt to environments [69]. Finally, we confirmed pyruvate fermentation to acetoin and hydroxy-proline degradation for the host-associated microbiome, which we previously found to be associated with bacterial load and worm fitness and which could potentially be involved in worm attraction and skin microbiome interactions [13]. Taken together, we identified several functions potentially critical for *C. elegans* physiology to be enriched in the worm. These findings may suggest that this nematode specifically selects bacteria providing these functions. To date, it is unclear how exactly *C. elegans* is capable of selecting specific microbial taxa.

### 5.4 Conclusion

This work introduced an approach for studying changes in microbiota-mediated functions by combining microbial abundance data with genome-inferred metabolic network models. We employed metabolic modeling, pathway analysis, and ecological concepts to infer functions potentially relevant to the microbiome of *C. elegans*. Our time series data of a defined microbial community revealed that the host microbiome is shaped by selective rather than stochastic processes. These selective processes appear to favor an enrichment of microbiome-mediated functions in *C. elegans* that are typical for gut microbiomes and that are likely beneficial for the nematode.

## 6 Acknowledgement

We thank the Kaleta, Traulsen and Schulenburg groups and also Alejandra Zárate-Potes, Alexandre Benedetto, and Peter Deines for valuable advice on our project. We are grateful for funding by the German Science Foundation (Deutsche Forschungsgemeinschaft, DFG) wthin the Collaborative Research Center 1182 (Project-ID 261376515), projects A1.1 (to HS), A1.5 (CK), A4.1 (AT), A4.3 (HS), and INF (LMS). We further acknowledge financial support by the Max-Planck Society (institutional funding to AT, fellowship to HS).

## References

[1] Margaret McFall-Ngai et al. “Animals in a bacterial world, a new imperative for the life sciences.” eng. In: Proc Natl Acad Sci U S A 110.9 (Feb. 2013), pp. 3229–3236. url: http://dx.doi.org/10.1073/pnas.1218525110.

[2] Harry J. Flint et al. “Microbial degradation of complex carbohydrates in the gut.” In: Gut Microbes 3.4 (2012), pp. 289–306. url: http://dx.doi.org/10.4161/gmic.19897.

[3] Roeland C H J van Ham et al. “Reductive genome evolution in Buchnera aphidicola.” In: Proc. Natl. Acad. Sci. U.S.A. 100 (2003), pp. 581–6. url: https://doi.org/10.1073/pnas.0235981100.

[4] Maria Gloria Dominguez-Bello et al. “Role of the microbiome in human development.” In: Gut 68 (2019), pp. 1108–1114. url: https://doi.org/10.1136/gutjnl-2018-317503.

[5] Eric Muraille. “Redefining the immune system as a social interface for cooperative processes.” eng. In: PLoS Pathog 9.3 (Mar. 2013), e1003203. url: http://dx.doi.org/10.1371/journal.ppat.1003203.

[6] Yael Litvak et al. “Commensal Enterobacteriaceae Protect against Salmonella Colonization through Oxygen Competition”. In: Cell host & microbe 25.1 (2019), pp. 128–139.

[7] Brittany D Needham et al. “A gut-derived metabolite alters brain activity and anxiety behaviour in mice”. en. In: Nature 602.7898 (Feb. 2022), pp. 647–653.

[8] Adam R Burns et al. “Contribution of neutral processes to the assembly of gut microbial communities in the zebrafish over host development”. In: The ISME journal 10.3 (2016), pp. 655–664.

[9] Mark Vellend. The theory of ecological communities. Monographs in population biology 57. Princeton, New Jersey: Princeton University Press, 2016. ix, 229 Seiten.

[10] Diana R. Nemergut et al. “Patterns and processes of microbial community assembly.” eng. In: Microbiol Mol Biol Rev 77.3 (Sept. 2013), pp. 342–356. url: http://dx.doi.org/10.1128/MMBR.00051-12.

[11] Philipp Dirksen et al. “The native microbiome of the nematode Caenorhabditis elegans: gateway to a new host-microbiome model”. In: BMC Biology 14.1 (May 2016).

[12] Julia Johnke, Philipp Dirksen, and Hinrich Schulenburg. “Community assembly of the native C. elegans microbiome is influenced by time, substrate and individual bacterial taxa.” In: Environ Microbiol 22 (2020), pp. 1265–1279. url: https://doi.org/10.1111/1462-2920.14932.

[13] Johannes Zimmermann* et al. “The functional repertoire contained within the native microbiota of the model nematode Caenorhabditis elegans.” In: ISME Journal 14.1 (2020), pp. 26–38. url: https://doi.org/10.1038/s41396-019-0504-y.

[14] Kohar A.B. Kissoyan et al. “Natural C. elegans Microbiota Protects against Infection via Production of a Cyclic Lipopeptide of the Viscosin Group”. In: Current Biology 29.6 (2019), 1030–1037.e5. url: http://www.sciencedirect.com/science/article/pii/S096098221930079X.

[15] Kohar Annie B Kissoyan et al. “Exploring Effects of C. elegans Protective Natural Microbiota on Host Physiology”. In: Frontiers in Cellular and Infection Microbiology 12 (2022), p. 53.

[16] Sirena Montalvo-Katz et al. “Association with soil bacteria enhances p38-dependent infection resistance in Caenorhabditis elegans”. In: Infect. Immun. 81.2 (Feb. 2013), pp. 514–520.

[17] Emma Watson et al. “Integration of metabolic and gene regulatory networks modulates the C. elegans dietary response”. en. In: Cell 153.1 (Mar. 2013), pp. 253–266.

[18] Emma Watson et al. “Interspecies systems biology uncovers metabolites affecting C. elegans gene expression and life history traits”. en. In: Cell 156.4 (Feb. 2014), pp. 759–770.

[19] Michael P O’Donnell et al. “A neurotransmitter produced by gut bacteria modulates host sensory behaviour”. en. In: Nature 583.7816 (July 2020), pp. 415–420.

[20] Michael Sieber et al. “Neutrality in the Metaorganism.” In: PLoS Biol. 17 (2019), e3000298. url: https://doi.org/10.1371/journal.pbio.3000298.

[21] Johannes Zimmermann, Christoph Kaleta, and Silvio Waschina. “gapseq: informed prediction of bacterial metabolic pathways and reconstruction of accurate metabolic models”. In: Genome Biology 22.1 (2021). url: https://doi.org/10.1186/s13059-021-02295-1.

[22] Philipp Dirksen et al. “CeMbio - The Caenorhabditis elegans Microbiome Resource”. In: G3: Genes, Genomes, Genetics (July 2020), g3.401309.2020. url: https://doi.org/10.1534/g3.120.401309.

[23] Anton Bankevich et al. “SPAdes: a new genome assembly algorithm and its applications to single-cell sequencing.” eng. In: J Comput Biol 19.5 (May 2012), pp. 455–477. url: http://dx.doi.org/10.1089/cmb.2012.0021.

[24] A. V. Zimin et al. “The MaSuRCA genome assembler”. In: Bioinformatics 29.21 (Nov. 2013), pp. 2669–2677. url: http://dx.doi.org/10.1093/bioinformatics/btt476.

[25] Ryan R Wick et al. “Unicycler: Resolving bacterial genome assemblies from short and long sequencing reads.” In: PLoS Comput. Biol. 13 (2017), e1005595. url: https://doi.org/10.1371/journal.pcbi. 1005595.

[26] Torsten Seemann. Shovill: Assemble bacterial isolate genomes from Illumina paired-end reads. github. 2020. url: https://github.com/tseemann/shovill.

[27] Alexandre Souvorov, Richa Agarwala, and David J Lipman. “SKESA: strategic k-mer extension for scrupulous assemblies”. en. In: Genome Biol. 19.1 (Oct. 2018), p. 153.

[28] Alla Mikheenko et al. “Versatile genome assembly evaluation with QUAST-LG.” In: Bioinformatics 34 (2018), pp. i142.#x2013;i150. url: https://doi.org/10.1093/bioinformatics/bty266.

[29] Thorsten Seemann. ABRicate: Mass screening of contigs for antimicrobial and virulence genes. github. 2020. url: https://github.com/tseemann/abricate.

[30] Lihong Chen et al. “VFDB 2016: hierarchical and refined dataset for big data analysis—10 years on”. In: Nucleic acids research 44.D1 (2015), pp. D694–D697. url: https://doi.org/10.1093/nar/gkv1239.

[31] Alessandra Carattoli et al. “In silico detection and typing of plasmids using PlasmidFinder and plasmid multilocus sequence typing”. en. In: Antimicrob. Agents Chemother. 58.7 (July 2014), pp. 3895–3903.

[32] Ea Zankari et al. “Identification of acquired antimicrobial resistance genes”. en. In: J. Antimicrob. Chemother. 67.11 (Nov. 2012), pp. 2640–2644.

[33] Enrique Doster et al. “MEGARes 2.0: a database for classification of antimicrobial drug, biocide and metal resistance determinants in metagenomic sequence data”. en. In: Nucleic Acids Res. 48.D1 (Jan. 2020), pp. D561–D569.

[34] Victòria Pascal Andreu et al. “The gutSMASH web server: automated identification of primary metabolic gene clusters from the gut microbiota”. In: Nucleic Acids Research 49.W1 (May 2021), W263–W270. eprint: https://academic.oup.com/nar/article-pdf/49/W1/W263/38842391/gkab353.pdf. url: https://doi.org/10.1093/nar/gkab353.

[35] Han Zhang et al. “dbCAN2: a meta server for automated carbohydrate-active enzyme annotation”. In: Nucleic Acids Research 46.W1 (2018), W95–W101. url: http://dx.doi.org/10.1093/nar/gky418.

[36] Eugen Bauer et al. “BacArena: Individual-based metabolic modeling of heterogeneous microbes in complex communities”. In: PLOS Computational Biology 13.5 (2017), pp. 1–22. url: https://doi.org/10.1371/journal.pcbi.1005544.

[37] Daniel Straub et al. “Interpretations of environmental microbial community studies are biased by the selected 16S rRNA (gene) amplicon sequencing pipeline”. en. In: Front. Microbiol. 11 (Oct. 2020), p. 550420.

[38] Benjamin J Callahan et al. “DADA2: High-resolution sample inference from Illumina amplicon data”. en. In: Nat. Methods 13.7 (July 2016), pp. 581–583.

[39] Paul J McMurdie and Susan Holmes. “phyloseq: an R package for reproducible interactive analysis and graphics of microbiome census data”. en. In: PLoS One 8.4 (Apr. 2013), e61217.

[40] Michael I. Love, Wolfgang Huber, and Simon Anders. “Moderated estimation of fold change and dispersion for RNA-seq data with DESeq2”. In: Genome Biology 15.12 (Dec. 2014), p. 550. url: https://doi.org/10.1186/s13059-014-0550-8.

[41] Zachary S L Foster, Thomas J Sharpton, and Niklaus J Grünwald. “Metacoder: An R package for visualization and manipulation of community taxonomic diversity data”. en. In: PLoS Comput. Biol. 13.2 (Feb. 2017), e1005404.

[42] William T Sloan et al. “Quantifying the roles of immigration and chance in shaping prokaryote community structure”. In: Environmental Microbiology 8.4 (2006), pp. 732–740.

[43] Daliang Ning et al. “A general framework for quantitatively assessing ecological stochasticity”. In: Proceedings of the National Academy of Sciences 116.34 (Aug. 2019), pp. 16892–16898. url: https://doi.org/10.1073/pnas.1904623116.

[44] Peter J Turnbaugh et al. “A core gut microbiome in obese and lean twins”. In: nature 457.7228 (2009), p. 480.

[45] J. Philip Grime and Simon Pierce. The evolutionary strategies that shape ecosystems. Wiley-Blackwell, 2012.

[46] Antonio Tamburro et al. “Expression of glutathione S-transferase and peptide methionine sulphoxide reductase in Ochrobactrum anthropi is correlated to the production of reactive oxygen species caused by aromatic substrates”. In: FEMS Microbiol. Lett. 241.2 (Dec. 2004), pp. 151–156.

[47] Wentao Yang et al. “The Inducible Response of the Nematode Caenorhabditis elegans to Members of Its Natural Microbiota Across Development and Adult Life”. In: Frontiers in Microbiology 10 (2019), p. 1793. url: https://www.frontiersin.org/article/10.3389/fmicb.2019.01793.

[48] Fan Zhang et al. “Natural genetic variation drives microbiome selection in the Caenorhabditis elegans gut”. In: Current Biology 31.12 (2021), 2603–2618.e9. url: https://www.sciencedirect.com/science/ article/pii/S0960982221005935.

[49] Maureen Berg, Xiao Ying Zhou, and Michael Shapira. “Host-Specific Functional Significance of Caenorhabditis Gut Commensals”. In: Frontiers in Microbiology 7 (Oct. 2016).

[50] Junjie Qin et al. “A metagenome-wide association study of gut microbiota in type 2 diabetes”. In: Nature 490.7418 (Oct. 2012), pp. 55–60.

[51] Johannes Mansfeld et al. “Branched-chain amino acid catabolism is a conserved regulator of physiological ageing”. In: Nat. Commun. 6.1 (Dec. 2015), p. 10043.

[52] Fan Zhang et al. “Caenorhabditis elegans as a Model for Microbiome Research”. In: Frontiers in Microbiology 8 (Mar. 2017). url: https://doi.org/10.3389%2Ffmicb.2017.00485.

[53] Alexander T. Neu, Eric E. Allen, and Kaustuv Roy. “Defining and quantifying the core microbiome: Challenges and prospects”. In: Proceedings of the National Academy of Sciences 118.51 (Dec. 2021).

[54] W. Ford Doolittle and S. Andrew Inkpen. “Processes and patterns of interaction as units of selection: An introduction to ITSNTS thinking”. In: Proceedings of the National Academy of Sciences 115.16 (Mar. 2018), pp. 4006–4014. url: https://doi.org/10.1073/pnas.1722232115.

[55] Jizhong Zhou and Daliang Ning. “Stochastic Community Assembly: Does It Matter in Microbial Ecology?” In: Microbiology and Molecular Biology Reviews 81.4 (2017), e00002–17. url: https://journals.asm.org/doi/abs/10.1128/MMBR.00002-17.

[56] David K Chow et al. “Sarcopenia in the Caenorhabditis elegans pharynx correlates with muscle contraction rate over lifespan”. en. In: Exp. Gerontol. 41.3 (Mar. 2006), pp. 252–260.

[57] Matthew A Jackson et al. “Signatures of early frailty in the gut microbiota”. en. In: Genome Med. 8.1 (Jan. 2016), p. 8.

[58] Tomasz Wilmanski et al. “Gut microbiome pattern reflects healthy ageing and predicts survival in humans”. In: Nature Metabolism 3.2 (Feb. 2021), pp. 274–286. url: https://doi.org/10.1038/s42255-021-00348-0.

[59] Victòria Pascal Andreu et al. “gutSMASH predicts specialized primary metabolic pathways from the human gut microbiota”. In: Nat. Biotechnol. (Feb. 2023).

[60] Bart van der Hee and Jerry M Wells. “Microbial regulation of host physiology by short-chain fatty acids”. In: Trends in Microbiology 29.8 (2021), pp. 700–712.

[61] Alyssa C. Walker et al. “Colonization of the Caenorhabditis elegans gut with human enteric bacterial pathogens leads to proteostasis disruption that is rescued by butyrate”. In: 17.5 (May 2021). Ed. by Andreas J. Baumler, e1009510. url: https://doi.org/10.1371/journal.ppat.1009510.

[62] Emma Watson et al. “Metabolic network rewiring of propionate flux compensates vitamin B12 deficiency in C. elegans”. In: eLife 5 (July 2016). url: https://doi.org/10.7554/eLife.17670.

[63] Snehal N Chaudhari et al. “Bacterial folates provide an exogenous signal for C. elegans germline stem cell proliferation”. In: Developmental cell 38.1 (2016), pp. 33–46.

[64] Bhupinder Virk et al. “Folate Acts in E. coli to Accelerate C. elegans Aging Independently of Bacterial Biosynthesis”. In: Cell Reports 14.7 (2016), pp. 1611–1620. url: https://www.sciencedirect.com/science/article/pii/S2211124716300298.

[65] Gunnar C Hansson. “Mucins and the Microbiome.” In: Annu. Rev. Biochem. 89 (2020), pp. 769–793. url: https://doi.org/10.1146/annurev-biochem-011520-105053.

[66] Antony Bacic, Itzhak Kahane, and Bert M Zuckerman. “Panagrellus redivivus and Caenorhabditis elegans: evidence for the absence of sialic acids”. In: Experimental parasitology 71.4 (1990), pp. 483–488.

[67] Roy G Cutler et al. “Sphingolipid metabolism regulates development and lifespan in Caenorhabditis elegans”. en. In: Mech. Ageing Dev. 143-144 (Dec. 2014), pp. 9–18.

[68] Ronald L. Schnaar. “The Biology of Gangliosides”. In: Sialic Acids, Part II: Biological and Biomedical Aspects. Ed. by David C. Baker. Vol. 76. Advances in Carbohydrate Chemistry and Biochemistry. Academic Press, 2019, pp. 113–148. url: https://www.sciencedirect.com/science/article/pii/S0065231818300027.

[69] Davide Arella, Maddalena Dilucca, and Andrea Giansanti. “Codon usage bias and environmental adaptation in microbial organisms”. en. In: Mol. Genet. Genomics 296.3 (May 2021), pp. 751–762.

