## Supplementary material for "Gut-associated functions are favored during microbiome assembly across *C. elegans* life": Supplemantary Information

March 23, 2023

2 Isolation and Inoculation details 6

3 Genome assembly 7

4 Quality 16S amplicon sequencing 10

6 Abundances 13

7 $\alpha$-diversity 14

8 $\beta$-diversity 16

9 Stochasticity 19

10 Functional abundances 20

11 Software 29

### 1 Extended material and methods

#### 1.1 Material

##### 1.1.1 Nematode strain and maintenance

The *C. elegans* strain DH26 (obtained from the Caenorhabditis Genetics Center; CGC, Minnesota, USA) was used, because it is spermatogenesis defective and thus sterile at 25 °C, avoiding overlapping generations in our experiment. This strain was grown and maintained according to the standard maintenance protocol using nematode growth medium (NGM) plates seeded with *Escherichia coli* strain OP50 at 20 °C [30] Worm populations were developmentally synchronized by a bleaching protocol that simultaneously removes any associated microbes [30], leading to germfree first instar larvae (L1) for the experiment.

##### 1.1.2 Bacteria and maintenance

We defined a community of 43 natural *C. elegans* microbiome isolates, the CeMbio43 (Table 1, Table S1), which extends the previously described *Caenorhabditis elegans* microbiome resource (CeMbio) that consists of 12 bacterial strains, representative of the naturally associated microbiome of this nematode [8]. The CeMbio43 community now includes additional taxa which we recently found to be commonly associated with *C. elegans* in nature [9, 14] and which also contains several strains for each of the abundant genera, in order to take account of strain redundancy commonly found in microbiomes. Thus, the CeMbio43 includes all 12 strains of the original CeMbio resource [8], five strains characterized previously by us in a different study [37], and another 26 strains isolated from natural *C. elegans* isolates or *C. elegans*-containing compost substrate, both collected in the Kiel Botanical Garden, Germany (Table 1, Table S1). This extended community earlier contained three more strains of the genera *Stenotrophomonas* (2x) and *Chryseobacterium* (1x), which we subsequently ignored because of cultivating problems and/or a potential pathogenic effect on worms. For DNA isolation and our experiments, the 43 bacterial strains of the CeMbio43 community were individually grown in 4 ml lysogeny broth (LB), using 48-Well 6 ml polypropylene storage plates and an inoculation of 4 µl bacterial stock culture (due to slow growth, 10 µl of *Gluconobacter* strains MYb592, MYb595, MYb596), followed by incubation for 2 days at 25 °C on a circular vibrating shaker at 350 rpm. Individual isolates were frozen in stocks of filter-sterilized 10 % DMSO. Each stock was only used once.

#### 1.2 Whole genome sequence analysis of bacteria

##### 1.2.1 DNA isolation and DNA sequencing

For the 26 bacterial strains without genome information S2, we isolated high quality genomic DNA using a modified CTAB protocol, as described previously [37]. Thereafter, short-read sequencing was performed by NextSeq 500 using the NextSeq 500/550 High Output Kit v2.5 (300 Cycles), and long-read sequencing with the Pacbio Sequel-54349U.

##### 1.2.2 Processing of sequencing data and genome assembly

Raw reads from genome sequencing were filtered using fastq_illumina_filter 0.1, trimmed with PRINSEQ-lite 0.20.4 [24] and re-synchronized by repair.sh (from BBMap version 38.87) [3]. The genome assembly was done with several algorithms and piplelines for comparison: i) SPAdes v3.14.1 [1], ii) MaSuRCA 3.4.2 [36], iii) Unicycler v0.4.8 (normal and bold mode), iv) [33], v) shovill 1.1.0 [26], v) SKESA 2.4.0 [29]. In addition to

short reads, long reads were available for some isolates (MYb115, MYb45, MYb264). For these cases, additional hybrid assemblies were obtained from SPAdes, MaSuRCA, and Unicycler.
The quality of the assembled genomes were evaluated using QUAST v5.1.0rc1 [20]. For each assembly, a score was calculated based on characteristics such as number of contigs, N50, L50, largest contig, completeness, mapped reads, properly paired reads, and N's per 100 kbp. The score was increased by 1 if the value of a characteristic belonged to the best 25% of all assemblies for an organisms. The characteristics 'completeness' and 'longest contig' were considered as most important and the score was increased by 2 if their value belonged to the best 5%. All assembly contigs were scrutinized for potential contamination using blobtools 1.1.1 [15] (Figure S1). For this means, short raw reads were sorted with samtools 1.10 [17] and mapped against the assembled contigs by BWA [16]. The assembled contigs were aligned against the nucleotide database (NCBI nt, 09/2019) employing blastn [5]. As a consequence, a contig classified by blobtools as Nematode was removed for MYb398. Finally, basic polishing was done based on coverage length GC plots as proposed by [11]. Contigs were filtered by legnth (>500bp), coverage (>5), and GC (<0.95) and the remaining contigs were considered as the final genomic sequence. Whenever possible, processes were parallized with gnu parallel [32]. Details on different parameters used for programs are listed in Table S12.

##### 1.2.3 Annotation of genomic functions

For all bacterial strains, we used the genome sequencing data to annotate genomic functions. In detail, the metabolic pathways were predicted, network models reconstructed, growth media inferred, and carbon source and fermentation products determined using gapseq 1.1 (Sequence DB: 1139b8e, bitscore cutoff 150). Virulence, resistance genes, and plasmids were scanned with abricate 1.0.1 [25] using the databases vfdb 2018-Jul-16 [7], plasmidfinder 2022-Mar-8 [6], resfinder 2022-Mar-8 [34], megares 2022-Mar-8 [10]. Gut microbiome specific gene clusters were identified by gutSMASH 1.0.0-1555cd7 [23]. For CAzyme annotation, dbCAN 3.0.2 was used [35]. The microbial interactions were simulated by a pairwise comparison of single vs. co-growth rate assuming TSB growth medium with BacArena 1.8.2 [2]. Details on different parameters used for programs are listed in Table S12.

#### 1.3 Analysis of microbiome changes

##### 1.3.1 Experiments and processing of samples

Temporal changes in microbiota composition were assessed for nematode populations, the directly connected NGM plate environments (labelled 'substrate'), and the control NGM plates without worms (labelled 'control') in ten replicates and across six time points. DH26 *C. elegans* populations were synchronized directly before start of the experiment. The bacterial inoculum was prepared by mixing 200 to 2000 µl of individual bacterial cultures, ensuring similar cell numbers per strain (Table S1). The inoculum was washed three times with sterile phosphate-buffered saline (PBS), followed by inoculation of NGM plates with 500 µl of the mix at OD(600nm) 1 (t0) and subsequent incubation for two hours at 25 °C (t1). Samples from t0 and t1 were frozen for later analysis. Around 500 L1 nematodes were added per plate and maintained at 25 °C. Nematodes and bacterial lawns were sampled after 16 h (t2), 42 h (t3), 66 h (t4), 90 h (t5), 138 h (t6), and 186 h (t7).
At each time point, nematodes and bacterial lawns were washed off the plates. Worms were separated from environmental microbiota using a modification of a previously published washing protocol [22]. In detail, 2 ml of each sample was distributed over the tops of five sterile filter tips (PreCision SafeSeal LC2695, Biozym) placed on specifically designed autoclaved aluminium boxes [22]. The filter tips were sealed with aluminium foil to prevent spill-over and cross-contamination. The boxes were centrifuged at 3000 rpm for 1 min, permitting isolation of all *C. elegans* developmental stages from the filters of the tips, while the wash-through including the environmental bacteria is collected in the wells of the aluminium box. The wash-through of each individual sample was then transferred to a sterile 1.5 ml tube. The tubes were centrifuged at 5000 rpm for 10 min, the supernatant was discarded, and the pellet was resuspended in 195 µl 5xGoTaq buffer (Promega, Madison, USA)

and 5 µl Proteinase K and frozen at -80 °C until further use.
The worms from each individual sample were collected from the filters by resuspension with 250 µL filter-sterilized M9-T (M9-buffer incl. 0.025% Triton X-100) and collected in a sterile 1.5 ml reaction tube. The worms were allowed to settle, followed by transfer of 100 µl worm pellet to a new tube including 100 µl 10 mM tetramisole hydrochloride (Sigma-Aldrich Chemie GmbH, Steinheim, Germany) in M9-T. Tetramisole paralyzes the worms and thus prevents loss of gut-associated bacteria. Any surface adherent bacteria were removed by incubating the worms with 200 µL 2% alkaline hypochloride solution in M9-T for 2 min [8]. We checked the tubes individually to avoid disruption of the worm tissue. Approximately 300 µl of the supernatant was removed and the samples were washed five times with 1 ml filter-sterilized PBS-T (PBS incl. 0.025 % Triton X). The number of worms per tube was determined and the worms were transferred to a new tube containing 17 autoclaved 1 mm zirconium beads, topped up to 400 µl with PBS-T and disrupted for 3 min at 30 Hz (Bead Ruptor 96; Omni International, Inc. Kennesaw, USA). Samples were centrifuged at 5000 rpm for 10 min, the supernatant was discarded, and the pellet was resuspended in 195 µL 5xGoTaq buffer and 5 µL Proteinase K and frozen at -80 °C until further use.

##### 1.3.2 DNA isolation and 16S rDNA amplicon sequencing

DNA was extracted with the Nucleo Spin® 96 Tissue Kit (Macherey-Nagel) with the following modifications: After thawing the samples, 50 µl of each sample were mixed with 170 µl Lysis buffer (150 µl Buffer T1, 20 µl Proteinase K) and disrupted for 3 min at 30 Hz using thw Bead Ruptor 96. Following incubation at 56 °C overnight, 300 µl lysate were transferred to the wells of a RW block and mixed with 300 µl buffer BQ1 and 300 µl 99 % ethanol. Mock community dilution series (ZymoBIOMICS Microbial Community Standard; Catalog No. D6300)) and negative controls were included on each plate. The 16S library was prepared using the 341F (5'-CCTACGGGNGGCWGCAG-3') and 806R primer (5'-GACTACHVGGGTATCTAATCC-3') covering the V3–V4 region of the 16S rRNA gene. Libraries were sequenced on the Miseq platform using the MiSeq Reagent Kit 2 × 300 bp (Illumina).

##### 1.3.3 Analysis of microbial census data

We processed the obtained 16S amplicon sequencing data using the standardized amplicon sequencing pipeline nf-core/ampliseq 2.1.1 [31]. Sample sequences were inferred by DADA2 [4] using a truncation length of 280/230 for forward and reverse reads. Species were initially assigned employing the SILVA database (v138.1) for filtering, and afterwards, filtered sequences were assigned by alignment of reference 16S sequences for the CeMbio43 community inferred from the genomes (blastn -perc_identity 99 -qcov_hsp_perc 95).

The microbial composition data (ASV and taxonomic table) was analyzed with phyloseq 1.38.0 [19]. Low biomass samples are prone to index switching, a technical bias that hampers the interpretation of those samples [13]. When we compared the frequency of read counts per sample, we found a clear separation between samples with a low number of reads and most samples with a high number of reads, with the low-reads samples being evenly spread across sample types (Figure S4). Therefore, to minimize any risk of artifacts, we removed samples with less than 12,0000 reads. ASVs with uncharacterized phylum or ASVs belonging to phyla with low prevalence ($< 10$) were also removed, which was also the threshold below which Cyanobacteria occurred (Table S4). Various ASVs were agglomerated by aligning the inferred 16S sequence against the 16S references data using blastn and summing up all matched ASVs to now 33 ASVs that correspond to the observed members of the CeMbio community. Some strains of the CeMbio43 community showed very high similarities in the V3V4 region of their rRNA to one another, which rendered them indistinguishable for the 16S analysis (Table S3). These highly similar strains were on purpose part of the CeMbio43 community to acknowledge high redundancy of taxonomically and functionally highly similar strains, often found in microbiomes (Figure 1). For all cases, where strains could not be distinguished, we used combined strain names. ASVs that could not be matched to members of the CeMbio43 community were labeled as 'other' and kept during the analysis, therey accounting for a certain level of random noise. The alpha diversity was determined by the function `plot_richness` of phyloseq. DESeq2 1.34.0 was used to find differentially abundant species [18] that were visualized by heat trees using the R package metacoder 0.3.5.001 [12].

##### 1.3.4 Stochasticity

To assess the neutrality of the samples the expected long-term distribution of a neutral model was fitted to the sample abundance data [27, 28]. The goodness-of-fit measure $R^2$ then quantifies how well the abundance data conforms to a distribution expected from purely neutral dynamics. The Python script that was used is available at `www.github.com/misieber/neufit`. To evaluate the impact of stochastic processes on community assembly, a null model called Normalized Stochasticity Ratio (NST) was evaluated using the R package NST 3.1.10 [21] (function `tNST`. Sample source was used as treatment parameter (`treat.t`) and the null model with dirichlet distribution was used as it the default for relative or proportional abundance data. P-values for comparisons were determined by bootstrapping using the function `nst.boot` with 1000 randomizations (output variable `p.count`).

##### 1.3.5 Functional abundances

We combined the microbial 16S data with the functions predicted from genomic analysis and modeling (1.2.3). For each organism, the presence (true/false) of a particular function indicated if the organism's abundance contributed to the abundance of the function. Finally, by adding up the abundances of all species in a sample for which the presence of a function was predicted, the overall abundance of the function was determined. Like the species-level analysis, DESeq2 1.34.0 was then used to identify differentially abundant functions.

#### 2 Isolation and Inoculation details

| Species | ID | Lab origin | Source | Worm strain | Volume |
|---|---|---|---|---|---|
| *Acinetobacter guillouiae* | MYb10 | Schulenburg Lab, Kiel, Germany | worm | MY316 | 1000 µl |
| *Pseudomonas lurida* | MYb11 | Schulenburg Lab, Kiel, Germany | worm | MY316 | 2000 µl |
| *Comamonas sp. B-9* | MYb21 | Schulenburg Lab, Kiel, Germany | worm | MY316 | 2000 µl |
| *Chryseobacterium culicis* | MYb25 | Schulenburg Lab, Kiel, Germany | worm | MY316 | 500 µl |
| *Ochrobactrum anthropi* | MYb49 | Schulenburg Lab, Kiel, Germany | worm | MY316 | 500 µl |
| *Ochrobactrum pituitosum* | MYb58 | Schulenburg Lab, Kiel, Germany | worm | MY316 | 200 µl |
| *Comamonas sp. TK41* | MYb69 | Schulenburg Lab, Kiel, Germany | worm | MY316 | 500 µl |
| *Ochrobactrum vermis* | MYb71 | Schulenburg Lab, Kiel, Germany | worm | MY316 | 500 µl |
| *Pseudomonas fluorescence* | MYb115 | Schulenburg Lab, Kiel, Germany | worm | MY379 | 2000 µl |
| *Erwinia billingiae* | MYb121 | Schulenburg Lab, Kiel, Germany | worm | MY379 | 1000 µl |
| *Acinetobacter johnsonii* | MYb158 | Schulenburg Lab, Kiel, Germany | compost | NA | 2000 µl |
| *Enterobacter ludwigii* | MYb174 | Schulenburg Lab, Kiel, Germany | compost | NA | 500 µl |
| *Enterobacter ludwigii* | MYb176 | Schulenburg Lab, Kiel, Germany | compost | NA | 200 µl |
| *Acinetobacter sp. B2070* | MYb177 | Schulenburg Lab, Kiel, Germany | compost | NA | 2000 µl |
| *Sphingobacterium faecium* | MYb181 | Schulenburg Lab, Kiel, Germany | compost | NA | 500 µl |
| *Enterobacter sp. 638* | MYb186 | Schulenburg Lab, Kiel, Germany | compost | NA | 500 µl |
| *Acinetobacter sp. 4D-W-22* | MYb191 | Schulenburg Lab, Kiel, Germany | compost | NA | 2000 µl |
| *Enterobacter cloacae* | CEent1 | Shapira Lab, Berkeley, CA, US | worm | N2 | 200 µl |
| *Pseudomonas berkeleyensis* | MSPm1 | Shapira Lab, Berkeley, CA, US | worm | N2 | 2000 µl |
| *Chryseobacterium sp.* | MYb264 | Schulenburg Lab, Kiel, Germany | worm | MY2771 | 500 µl |
| *Chryseobacterium sp.* | MYb328 | Schulenburg Lab, Kiel, Germany | worm | MY2768 | 200 µl |
| *Pseudomonas sp.* | MYb330 | Schulenburg Lab, Kiel, Germany | worm | MY2768 | 200 µl |
| *Pseudomonas sp.* | MYb331 | Schulenburg Lab, Kiel, Germany | worm | MY2768 | 2000 µl |
| *Pseudomonas sp.* | MYb371 | Schulenburg Lab, Kiel, Germany | worm | MY2768 | 2000 µl |
| *Erwinia sp.* | MYb375 | Schulenburg Lab, Kiel, Germany | worm | MY2769 | 500 µl |
| *Ochrobactrum sp.* | MYb379 | Schulenburg Lab, Kiel, Germany | worm | MY2769 | 2000 µl |
| *Sphingobacterium sp.* | MYb382 | Schulenburg Lab, Kiel, Germany | worm | MY2769 | 200 µl |
| *Sphingobacterium sp.* | MYb388 | Schulenburg Lab, Kiel, Germany | worm | MY2769 | 1000 µl |
| *Comamonas sp.* | MYb396 | Schulenburg Lab, Kiel, Germany | worm | MY2769 | 2000 µl |
| *Pseudomonas sp.* | MYb398 | Schulenburg Lab, Kiel, Germany | worm | MY2769 | 200 µl |
| *Erwinia sp.* | MYb416 | Schulenburg Lab, Kiel, Germany | worm | MY2771 | 1000 µl |
| *Erwinia sp.* | MYb535 | Schulenburg Lab, Kiel, Germany | worm | MY2768 | 2000 µl |
| *Pseudomonas sp.* | MYb541 | Schulenburg Lab, Kiel, Germany | worm | MY2768 | 2000 µl |
| *Lelliottia amnigena* | JUb66 | Félix Lab, Paris, France | rotting apple | NA | 2000 µl |
| *Pantoea nemavictus* | BIGb0393 | Samuel Lab, Houston, TX, US | rotting petasites stem | NA | 2000 µl |
| *Sphingomonas molluscorum* | JUb134 | Félix Lab, Paris, France | worm | NA | 2000 µl |
| *Stenotrophomonas indicatrix* | JUb19 | Félix Lab, Paris, France | rotting pear | NA | 2000 µl |
| *Comamonas piscis* | BIGb0172 | Samuel Lab, Houston, TX, US | worm | NA | 200 µl |
| *Sphingobacterium multivorum* | BIGb0170 | Samuel Lab, Houston, TX, US | worm | NA | 200 µl |
| *Chryseobacterium scophthalmum* | JUb44 | Félix Lab, Paris, France | rotting apple | NA | 500 µl |
| *Gluconobacter wancherniae* | MYb592 | Schulenburg Lab, Kiel, Germany | rotting apple | NA | 2000 µl |
| *Gluconobacter cerinus* | MYb595 | Schulenburg Lab, Kiel, Germany | rotting apple | NA | 2000 µl |
| *Gluconobacter albidus* | MYb596 | Schulenburg Lab, Kiel, Germany | rotting apple | NA | 2000 µl |

Table S1: Isolation details for species included in the CeMbio43 bacterial community. Lab origin, the lab which originally isolated the bacterium. Source, isolation of the bacterium from either natural C. elegans or C. elegans-containing substrate. Worm strain, information on the specific C. elegans strain, from which the bacteria were isolated; NA, not applicable. Volume, volume of an overnight culture, which used to prepare the CeMbio mixture, which was used to inoculate the plates for the experiment.

### 3 Genome assembly

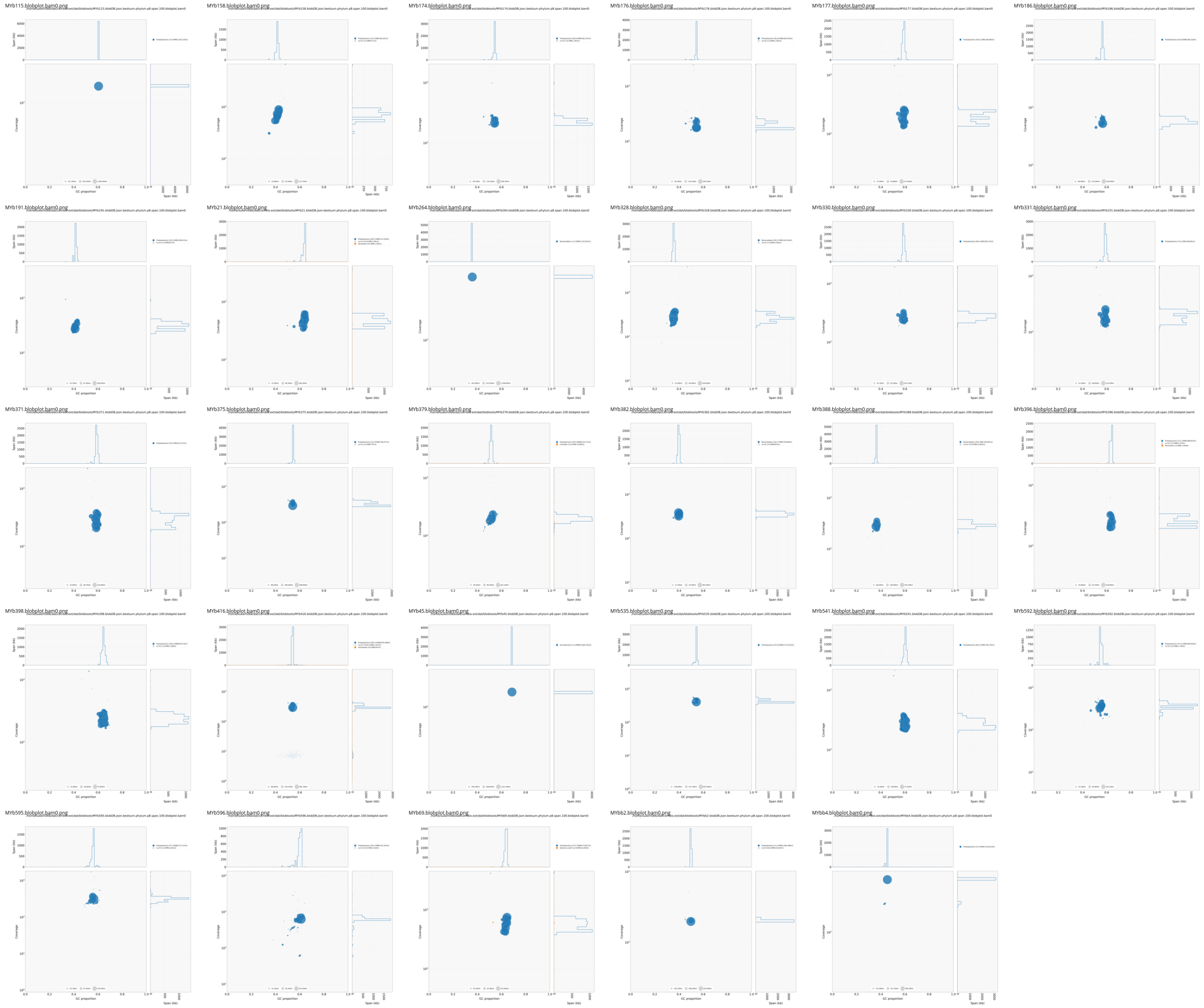

Figure S1: Contamination control of assembled contigs using Blobtools. Taxon-annotated GC-coverage plots were used to identify potential contaminations in genome assemblies.

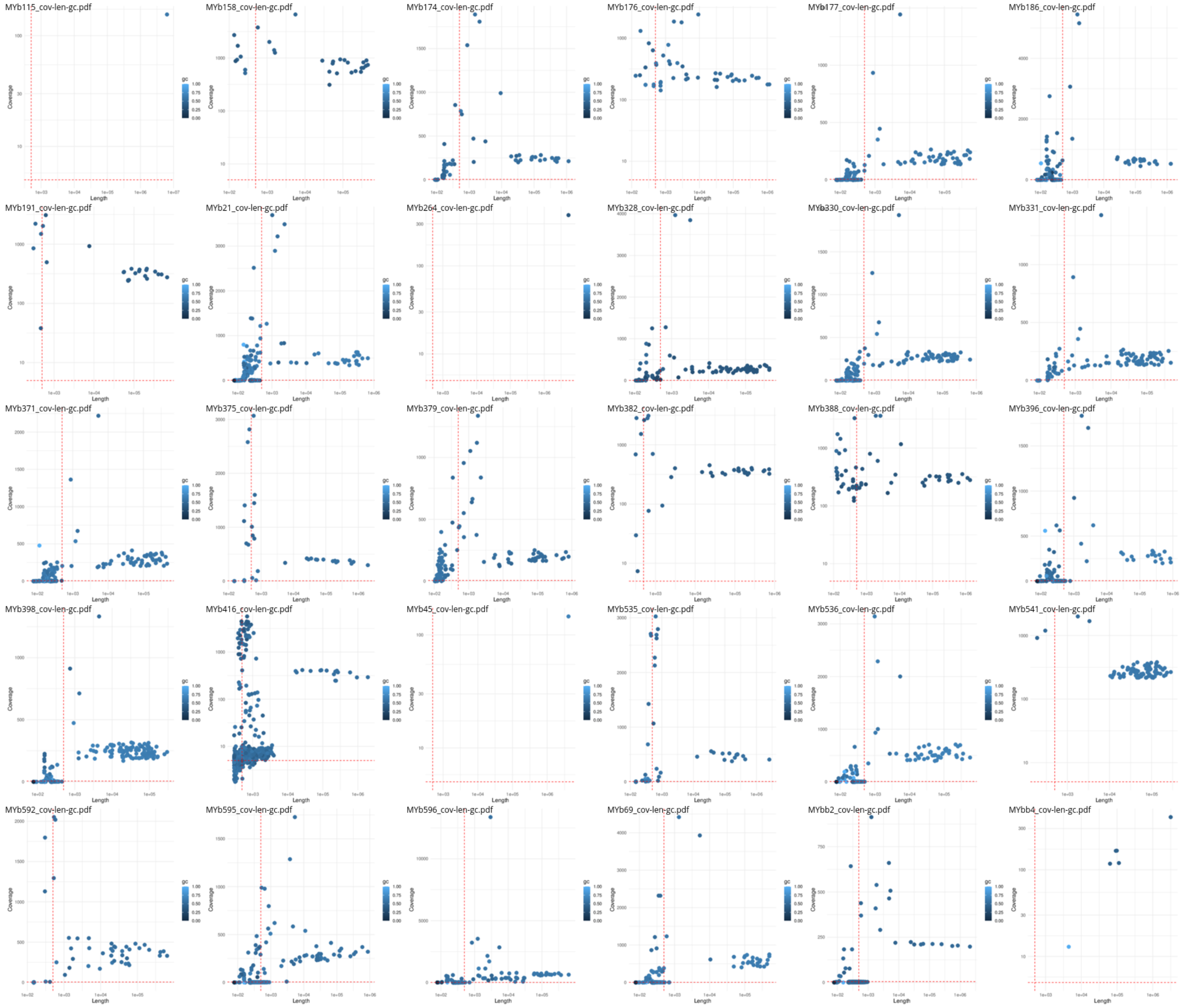

Figure S2: Coverage-length-GC plots of the assembled genomic contigs were used for basic polishing. We excluded contigs that had length < 500, coverage < 5, or GC>0.95 (red lines indicates these thresholds used for filtering).

| ID | Organism | Assembler | N50 | L50 | Contigs | Length | Completeness | Genes |
|---|---|---|---|---|---|---|---|---|
| MYb115 | *Pseudomonas fluorescence* | unicycler | 6342245 | 1 | 1 | 6342245 | 100.00 | 5767 |
| MYb158 | *Acinetobacter johnsonii* | unicycler | 346291 | 5 | 23 | 3599479 | 97.30 | 3425 |
| MYb174 | *Enterobacter ludwigii* | shovill | 482322 | 4 | 29 | 4642087 | 97.97 | 4325 |
| MYb176 | *Enterobacter ludwigii* | unicycler | 560529 | 3 | 37 | 4632577 | 97.97 | 4329 |
| MYb177 | *Acinetobacter sp. B2070* | spades | 266844 | 8 | 50 | 6107643 | 99.32 | 5563 |
| MYb186 | *Enterobacter sp. 638* | spades | 408218 | 4 | 26 | 4824469 | 98.65 | 4505 |
| MYb191 | *Acinetobacter sp. 4D-W-22* | masurca | 338615 | 4 | 21 | 3528846 | 96.62 | 3372 |
| MYb21 | *Comamonas sp. B-9* | spades | 312419 | 7 | 36 | 5363665 | 97.30 | 4802 |
| MYb264 | *Chryseobacterium sp.* | unicycler | 5234367 | 1 | 1 | 5234367 | 93.24 | 4562 |
| MYb328 | *Chryseobacterium sp.* | shovill | 163502 | 11 | 60 | 5570205 | 93.92 | 5006 |
| MYb330 | *Pseudomonas sp.* | spades | 206174 | 11 | 65 | 6101970 | 99.32 | 5563 |
| MYb331 | *Pseudomonas sp.* | shovill | 168801 | 12 | 71 | 6097985 | 99.32 | 5557 |
| MYb371 | *Pseudomonas sp.* | spades | 223371 | 10 | 53 | 6101404 | 99.32 | 5551 |
| MYb375 | *Erwinia sp.* | masurca | 706071 | 2 | 24 | 4907227 | 96.62 | 4512 |
| MYb379 | *Ochrobactrum sp.* | shovill | 524112 | 4 | 47 | 5072635 | 96.62 | 4855 |
| MYb382 | *Sphingobacterium sp.* | masurca | 530684 | 4 | 29 | 4738001 | 93.24 | 4023 |
| MYb388 | *Sphingobacterium sp.* | unicycler | 794901 | 3 | 29 | 6215939 | 93.24 | 5300 |
| MYb396 | *Comamonas sp.* | spades | 488653 | 4 | 23 | 5338904 | 97.30 | 4744 |
| MYb398 | *Pseudomonas sp.* | spades | 92388 | 21 | 105 | 5492602 | 100.00 | 5032 |
| MYb416 | *Erwinia sp.* | masurca | 601712 | 3 | 443 | 5789730 | 94.59 | 5710 |
| MYb535 | *Erwinia sp.* | masurca | 2570355 | 1 | 25 | 4921382 | 93.92 | 4604 |
| MYb541 | *Pseudomonas sp.* | unicycler | 109376 | 20 | 82 | 6347450 | 100.00 | 5874 |
| MYb592 | *Gluconobacter wancherniae* | masurca | 409404 | 4 | 40 | 3234488 | 95.27 | 3055 |
| MYb595 | *Gluconobacter cerinus* | spades | 371515 | 3 | 48 | 3591761 | 95.27 | 3345 |
| MYb596 | *Gluconobacter albidus* | spades | 191545 | 5 | 51 | 3298982 | 96.62 | 3068 |
| MYb69 | *Comamonas sp. TK41* | spades | 274857 | 7 | 28 | 5262490 | 97.30 | 4659 |

Table S2: Overview of newly assembled genomes. In total, 26 genomes were obtained using different assemblers. The program with the best assembly is shown together with the quality properties of the assembled genome.

### 4 Quality 16S amplicon sequencing

#### 4.1 Mock community

In Figure S3, the experimentally found composition of the Zymo mock community is compared with theoretical values provided by the manufacturer (ZymoBIOMICSTM Microbial Community Standard. Catalog No. D6300). *Salmonella* was only found at low abundances (<= 5%, category: 'x_other(low)') and *Lactobacillus* was classified as *Limosilactobacillus*. *Enterococcus* was found in two samples. One sample was removed because of a very low read count (< 100).

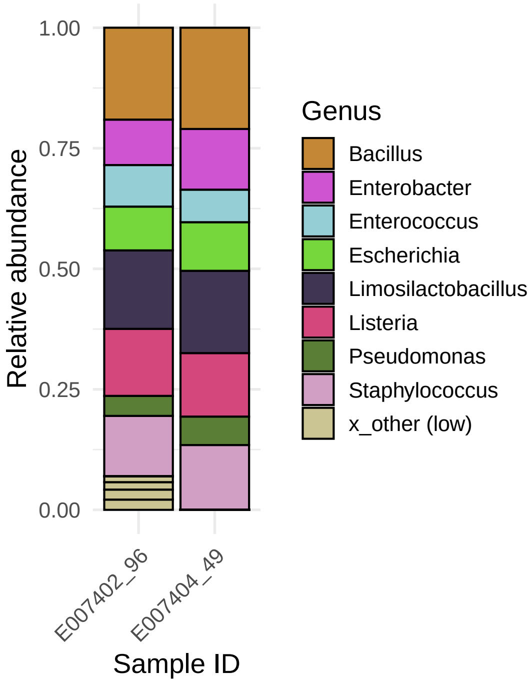

Figure S3: Zymo mock community: Experimentally found ASVs of a defined community was used for quality control. Label 'x_other (low)' indicates taxa with low maximal abundance $\leq 0.05$.

### 5 16S census data analysis

#### 5.1 Strains below detection limit

| Strain | Problem | Solution | Comment |
|---|---|---|---|
| MYb177 | genus unclear | ? | Acinetobacter (16S reference) vs. Pseudomonas (16S genome)? |
| MYb191, (MYb177) | same V3V4 | new name: MYb191* | matching V3V4 of MYb177 is from reference |
| MYb371, MYb331, MYb330, (MYb177) | same 16S | merge: MYb330-331-371 | genomic 16S identical, reference 16S differ, V3V4 for all identical; matching V3V4 of MYb177 is from genome |
| MYb396, MYb69, MYb21 | same V3V4 | merge: MYb21-69-396 | reference 16S differ, identical genomic 16S for MYb69 and MYb396 |
| MYb71, MYb49 | same V3V4 | merge: MYb49-71 | |
| MYb176, MYb174 | same V3V4 | merge: MYb174-176 | genomic 16 from rnammer quite distant to barrnap, reference close to barrnap (but same V3V4) |
| JUb66, MYb186 | same V3V4 | new name: MYb186* | JUb66 has unique hits that are also larger than combined hits |

Table S3: CeMbio43 strains below the detection limit. The table lists taxa for which the 16S amplicon sequencing could not detect certain strains. Problems arose because species could not be differentiated based on the V3V4 region. The table also mentioned what was done to address the problems.

#### 5.2 Filtering

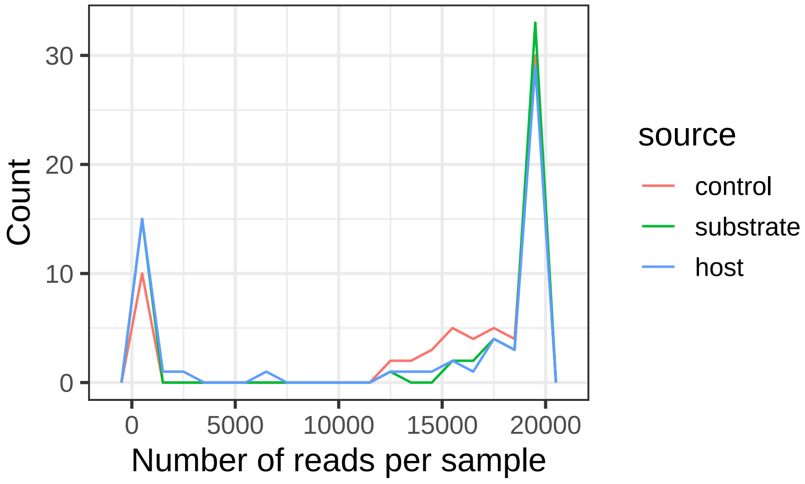

Figure S4: Frequency of samples with various read counts. All samples and their number of reads are shown. The figure displays a group of low-read-count samples and a group of high-read-count samples. Samples with 20000 or more reads were grouped.

| Phylum | Prevalence (mean) | Prevalence (sum) | Abundance |
|---|---|---|---|
| Proteobacteria | 12.5044 | 4239 | 0.7271 |
| Bacteroidetes | 103.6154 | 1347 | 0.2679 |
| Bacteroidota | 6.7720 | 1307 | 0.0048 |
| Firmicutes | 0.4189 | 31 | 0.0001 |
| Actinobacteriota | 0.6600 | 33 | 0.0001 |
| Cyanobacteria | 0.7000 | 7 | 0.0000 |
| Bdellovibrionota | 0.5000 | 1 | 0.0000 |
| Chloroflexi | 0.5000 | 1 | 0.0000 |
| Patescibacteria | 0.7143 | 5 | 0.0000 |
| Myxococcota | 1.0000 | 1 | 0.0000 |
| Verrucomicrobiota | 1.0000 | 1 | 0.0000 |
| Planctomycetota | 1.0000 | 1 | 0.0000 |
| Acidobacteriota | 0.7500 | 3 | 0.0000 |
| Spirochaetota | 0.0000 | 0 | 0.0000 |
| Nitrospirota | 1.0000 | 1 | 0.0000 |
| Deinococcota | 0.0000 | 0 | 0.0000 |
| Gemmatimonadota | 1.0000 | 1 | 0.0000 |

Table S4: Prevalence was compared on the phylum level to identify a threshold for filtering. We removed ASVs belonging to phyla with very low prevalence (prevalence sum $< 10$).

### 6 Abundances

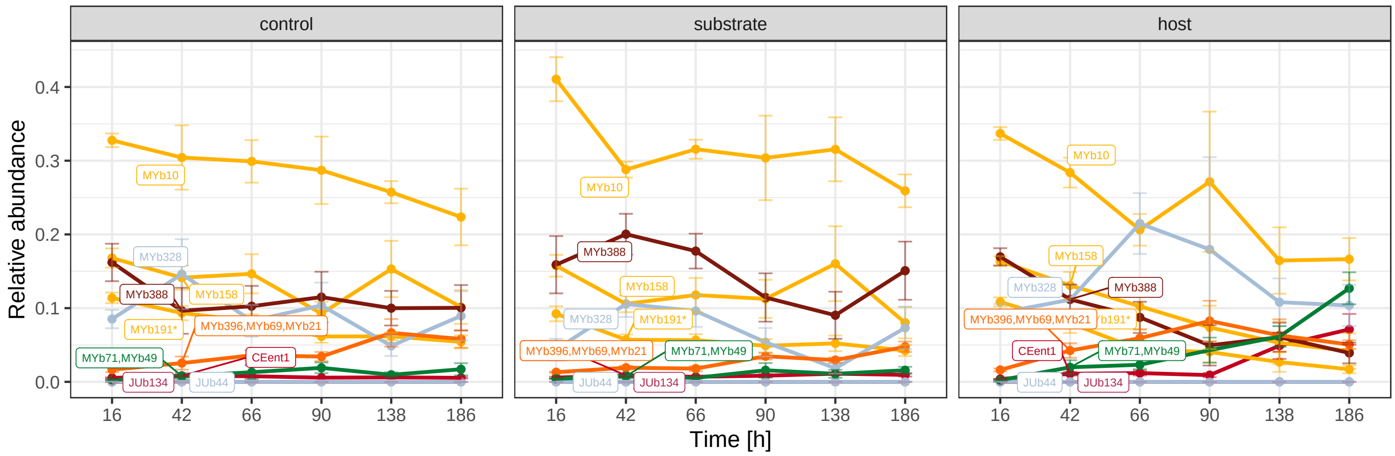

Figure S5: Relative abundances of taxa found to be differentially abundant between samples types (host, substrate, control). For each time point, mean relative abundances and standard errors are shown.

### 7 $\alpha$-diversity

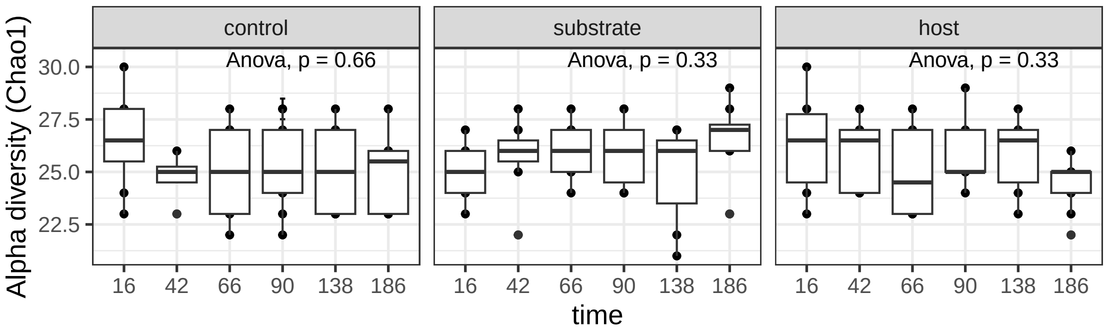

(a) Chao1

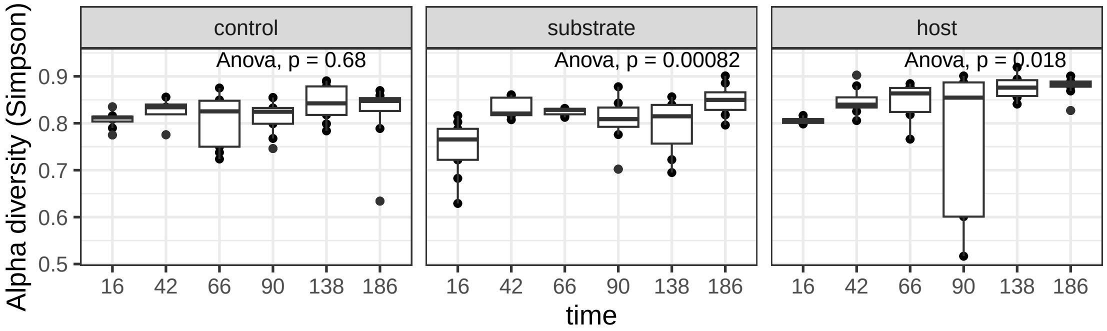

(b) Simpson

Figure S6: $\alpha$ diversity of filtered and agglomarated microbial samples on taxon level. Top row shows Richness (Chao1) and bottom row eveness (Simpson)

| | Df | Sum Sq | Mean Sq | F value | Pr(>F) |
|---|---|---|---|---|---|
| source | 2 | 0.6495 | 0.3248 | 6.640 | 0.0019 |
| time | 5 | 1.8774 | 0.3755 | 7.677 | 0.0000 |
| source:time | 10 | 0.7617 | 0.0762 | 1.557 | 0.1282 |
| Residuals | 116 | 5.6737 | 0.0489 | | |

Table S5: ANOVA test of Shannon $\alpha$-diversity checking for sample sources, time, and the interaction between sample source and time. Test statistics output from R is shown.

| time | diff | lwr | upr | p adj |
|---|---|---|---|---|
| 42-16 | 0.2766 | 0.0678 | 0.4853 | 0.0027 |
| 66-16 | 0.2484 | 0.0519 | 0.4449 | 0.0049 |
| 138-16 | 0.3186 | 0.1178 | 0.5194 | 0.0001 |
| 186-16 | 0.3737 | 0.1792 | 0.5683 | 0.0000 |
| 186-90 | 0.2036 | 0.0043 | 0.4029 | 0.0423 |

Table S6: Tukey Honest Significant Difference test for Shannon $\alpha$ diversity to check for differences between time points. Only significant pairs are show (p adj $< 0.05$). Test statistics output is obtained from R.

| source | diff | lwr | upr | p adj |
|---|---|---|---|---|
| substrate-control | -0.0158 | -0.1403 | 0.1088 | 0.9515 |
| host-control | 0.1417 | 0.0149 | 0.2685 | 0.0244 |
| host-substrate | 0.1575 | 0.0294 | 0.2856 | 0.0116 |

Table S7: Tukey Honest Significant Difference test for Shannon $\alpha$ diversity to check for differences between sample sources. The output of the test statistics from R is shown.

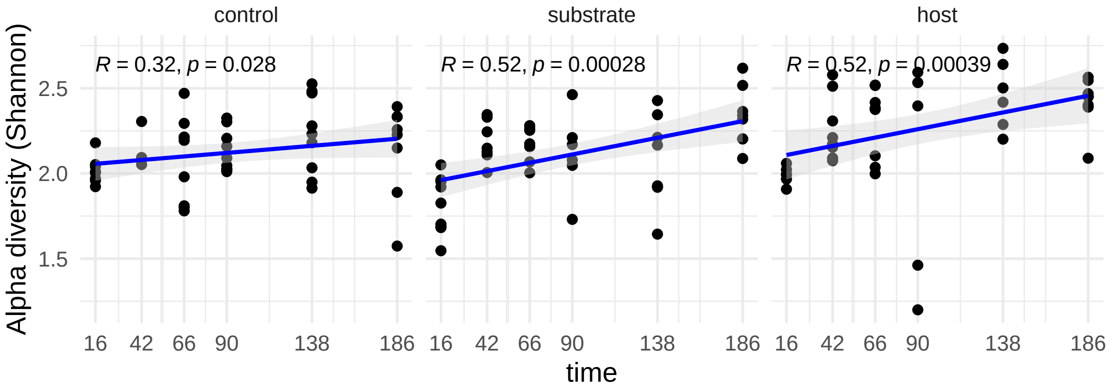

Figure S7: Shannon $\alpha$-diversity for sample sources is shown, including the linear regression (blue) and Spearman correlation test to visualize the diversity trend over time.

### 8 $\beta$-diversity

| | Df | SumOfSqs | R2 | F | Pr(>F) |
|---|---|---|---|---|---|
| source | 2 | 292.9 | 0.042 | 2.869 | 0.001 |
| Residual | 131 | 6686.4 | 0.958 | | |
| Total | 133 | 6979.3 | 1.000 | | |

Table S8: PERMANOVA test of differences between samples sources (Aitchison distance, adonis2 function of R packages vegan, 10000 permutations). The output of the test statistics from R is shown.

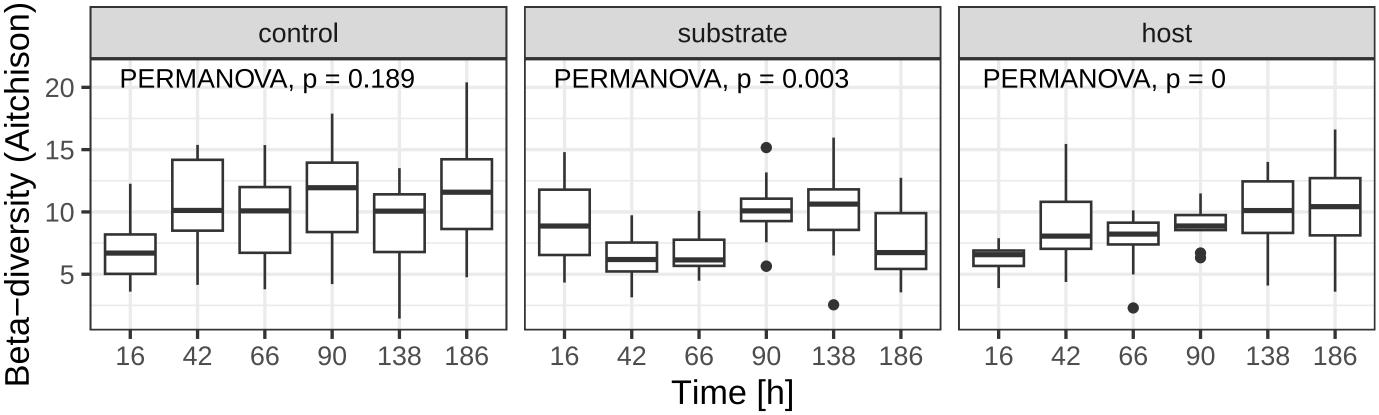

Figure S8: Time variation of the $\beta$-diversity for each sample type is shown. Aitchison distance was used to calculate dissimilarity. The significance of time variation was determined by a PERMANOVA test (adonis2 function of R packages vegan, 10000 permutations).

| time | dispersion test (bray) | permanova test (bray) | dispersion test (aitchison) | permanova test (aitchison) | dispersion test (unifrac) | permanova test (unifrac) |
|---|---|---|---|---|---|---|
| 16 | 0.1056 | 0.1028 | 0.1692 | 0.0589 | 0.2003 | 0.4251 |
| 42 | 0.7969 | 0.1536 | 0.4181 | 0.7873 | 0.6204 | 0.1691 |
| 66 | 0.2410 | 0.0092 | 0.1805 | 0.0700 | 0.1623 | 0.0080 |
| 90 | 0.8060 | 0.7112 | 0.2804 | 0.8509 | 0.6456 | 0.9090 |
| 138 | 0.7258 | 0.0202 | 0.9031 | 0.0855 | 0.7036 | 0.0507 |
| 186 | 0.3683 | 0.0018 | 0.2087 | 0.0108 | 0.7671 | 0.0091 |

Table S9: Dispersion test and PERMANOVA test of differences for each time point (Bray-Curtis, Aitchison, and weighted Unifrac distance; adonis2 and betadisper function of R package vegan, 10000 permutations). P-values for each test are shown.

| source | dispersion test (bray) | permanova test (bray) | dispersion test (aitchison) | permanova test (aitchison) | dispersion test (unifrac) | permanova test (unifrac) |
|---|---|---|---|---|---|---|
| control | 0.0193 | 0.3620 | 0.3078 | 0.1848 | 0.3060 | 0.6898 |
| substrate | 0.0279 | 0.0112 | 0.0975 | 0.0032 | 0.1418 | 0.0315 |
| host | 0.0059 | 0.0001 | 0.1412 | 0.0001 | 0.0074 | 0.0003 |

Table S10: Dispersion test and PERMANOVA test of differences for each sample source (Bray-Curtis, Aitchison, and weighted Unifrac distance; adonis2 and betadisper function of R package vegan, 10000 permutations). P-values for each test are shown.

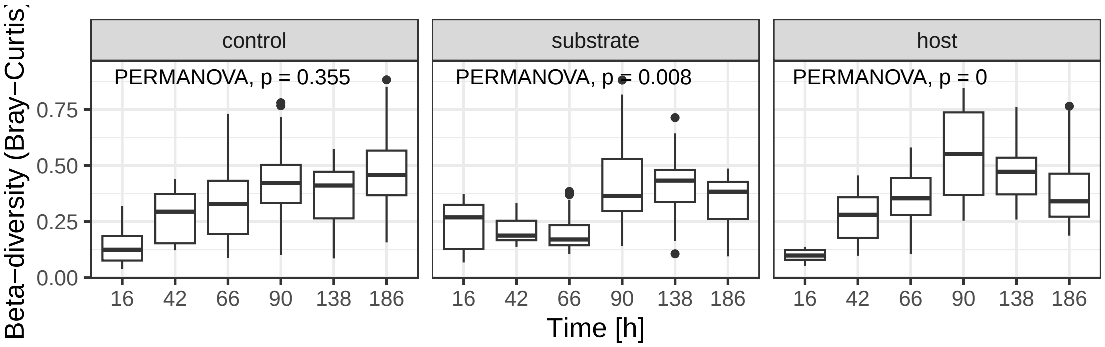

(a)

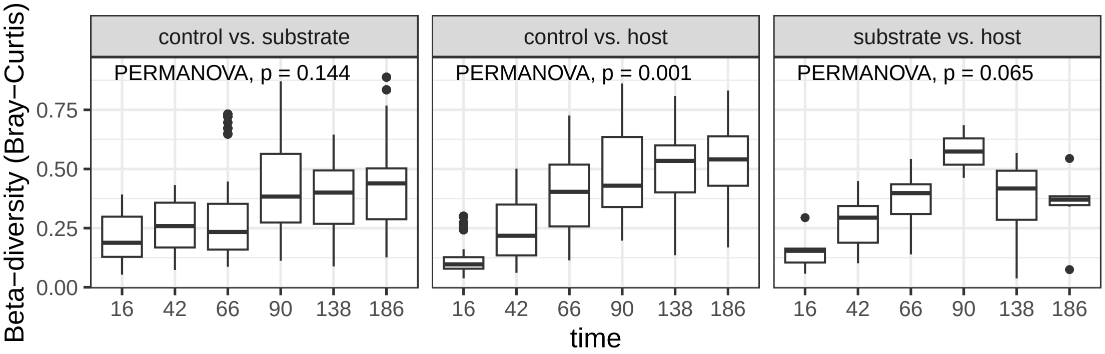

(b)

Figure S9: $\beta$-diversity using Bray-Curtis distance. Top row shows the results for each sample source over time. The bottom row shows differences compared between sample types. For comparing host-substrate samples, pairs from the same replicate were available, whereas for comparing host/control and substrate or control, pairs were randomly associated (100 repetitions), and the mean P-value is shown. The significance of time variation was determined by a PERMANOVA test (adonis2 function of R packages vegan, 10000 permutations).

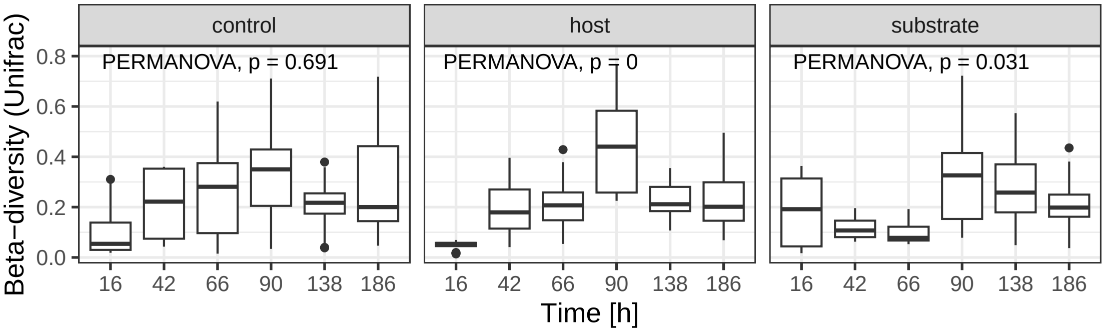

(a)

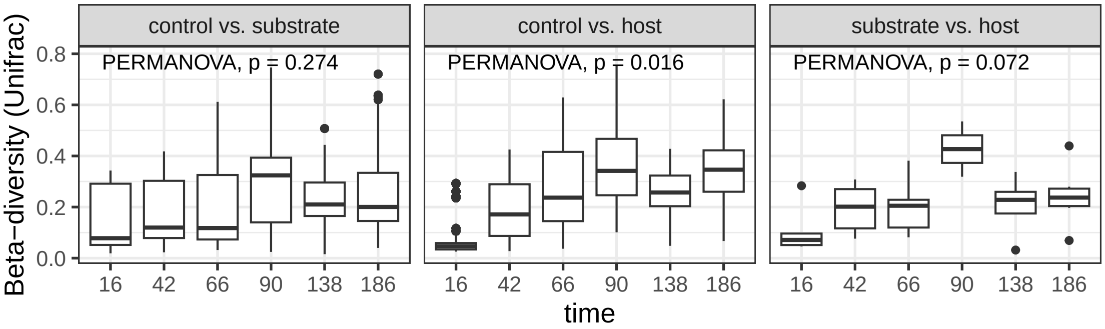

(b)

Figure S10: $\beta$-diversity using weighted Unifrac distance. Top row shows variation over time for each sample type. Bottom row shows differences compared between sample type. For comparing host-substrate samples, pairs from the same replicate were available, whereas for comparing host-control and substrate-control, pairs were randomly associated (100 repetitions), and the mean P-value is shown. The significance of time variation was determined by a PERMANOVA test (adonis2 function of R packages vegan, 10000 permutations).

### 9 Stochasticity

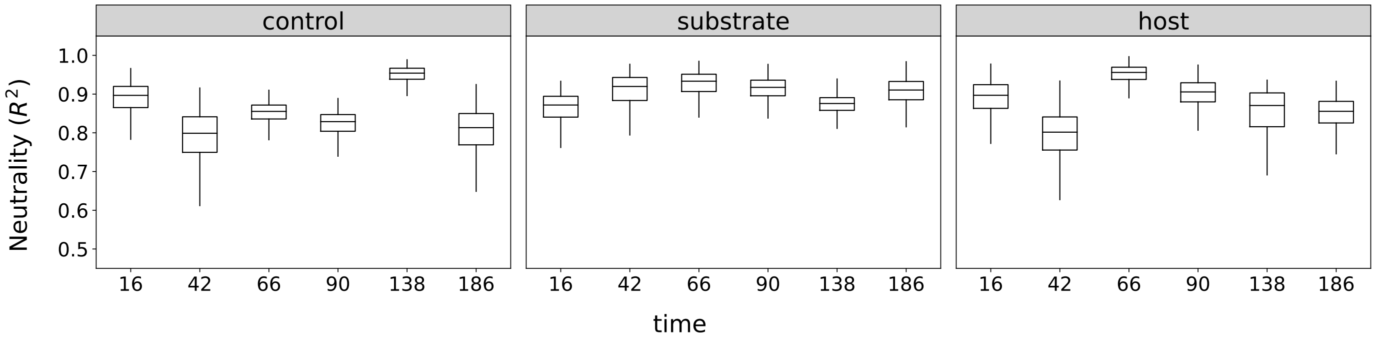

Figure S11: Goodness of fit of samples abundances to the neutral model. All samples sources are shown over time. High numbers of $R^2$ indicate that abundances match the neutral expectation.

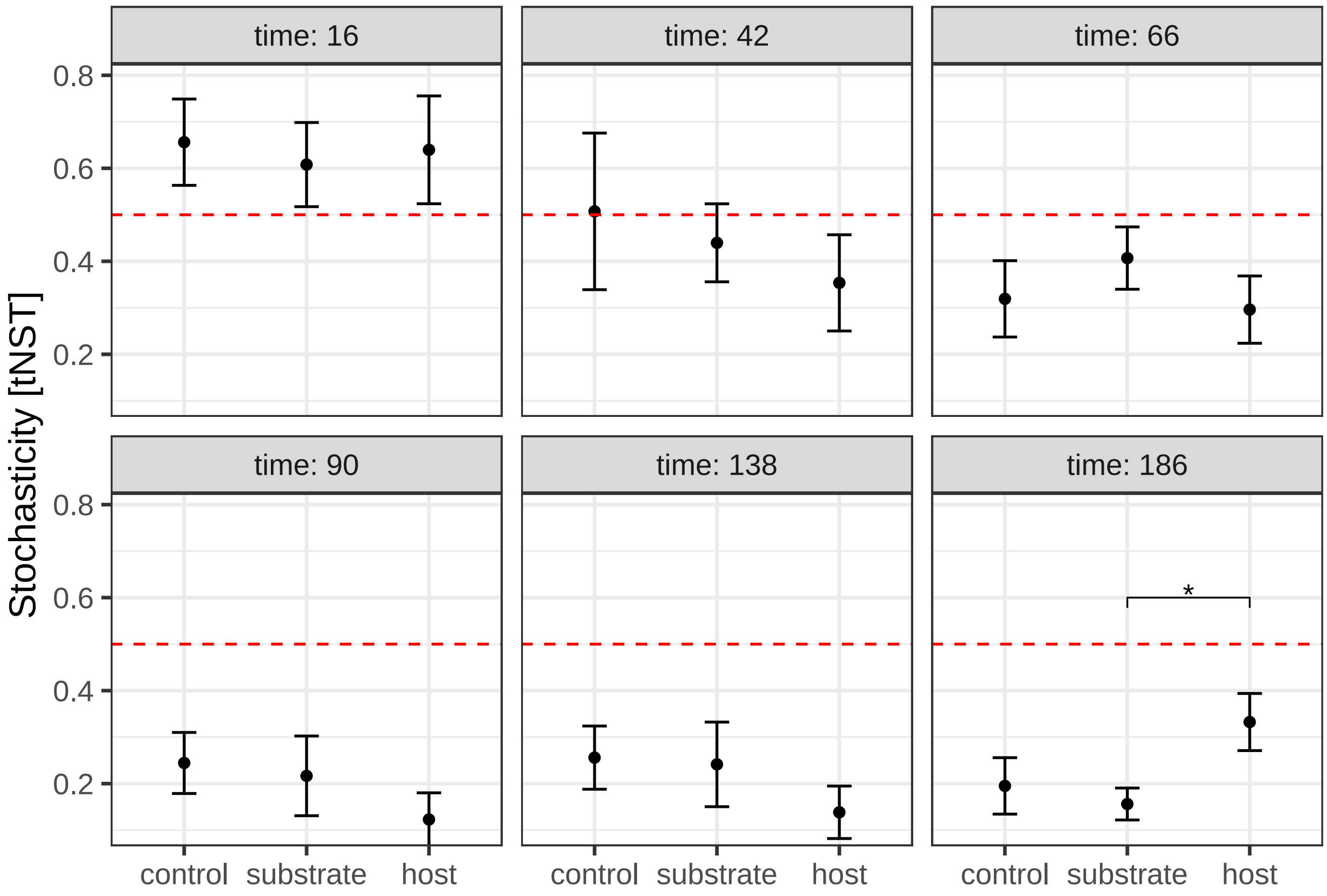

Figure S12: Taxonomic normalized stochasticity index (tNST). Comparison is shown between sample sources for every time point. The red line indicates the threshold between deterministic (tNST$< 0.5$) and stochastic (tNST$> 0.5$)community assembly.

#### 10 Functional abundances

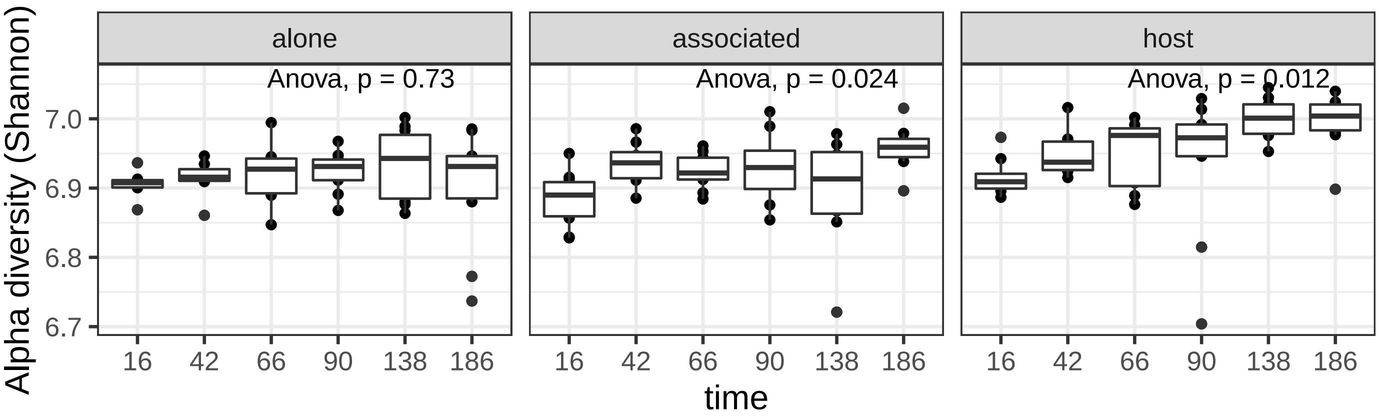

Figure S13: Shannon $\alpha$-diversity of predicted functions. Predicted functions were paired with species abundance data, and diversity analysis was performed on functional abundances similar to diversity analysis of species abundances.

| Comparison | Subsystem | GeneRatio | BgRatio | pvalue | p.adjust | qvalue | Count |
|---|---|---|---|---|---|---|---|
| host vs. substrate | Seconday metabolite degradation | 21/174 | 51/1005 | 0.0000 | 0.0007 | 0.0006 | 21 |
| substrate vs. host | Fatty acid and lipid degradation | 6/72 | 20/1005 | 0.0019 | 0.0343 | 0.0301 | 6 |
| host vs. control | Cofactor biosynthesis | 36/168 | 124/1005 | 0.0002 | 0.0037 | 0.0034 | 36 |
| control vs. host | Aromatic compound degradation | 25/117 | 84/1005 | 0.0000 | 0.0000 | 0.0000 | 25 |
| control vs. host | Amine degradation | 11/117 | 32/1005 | 0.0005 | 0.0054 | 0.0046 | 11 |

Table S11: Enrichment analysis of metabolic subsystems. We performed an over-representation analysis of differentially abundant metabolic pathways. The table shows significantly enriched subsystems.

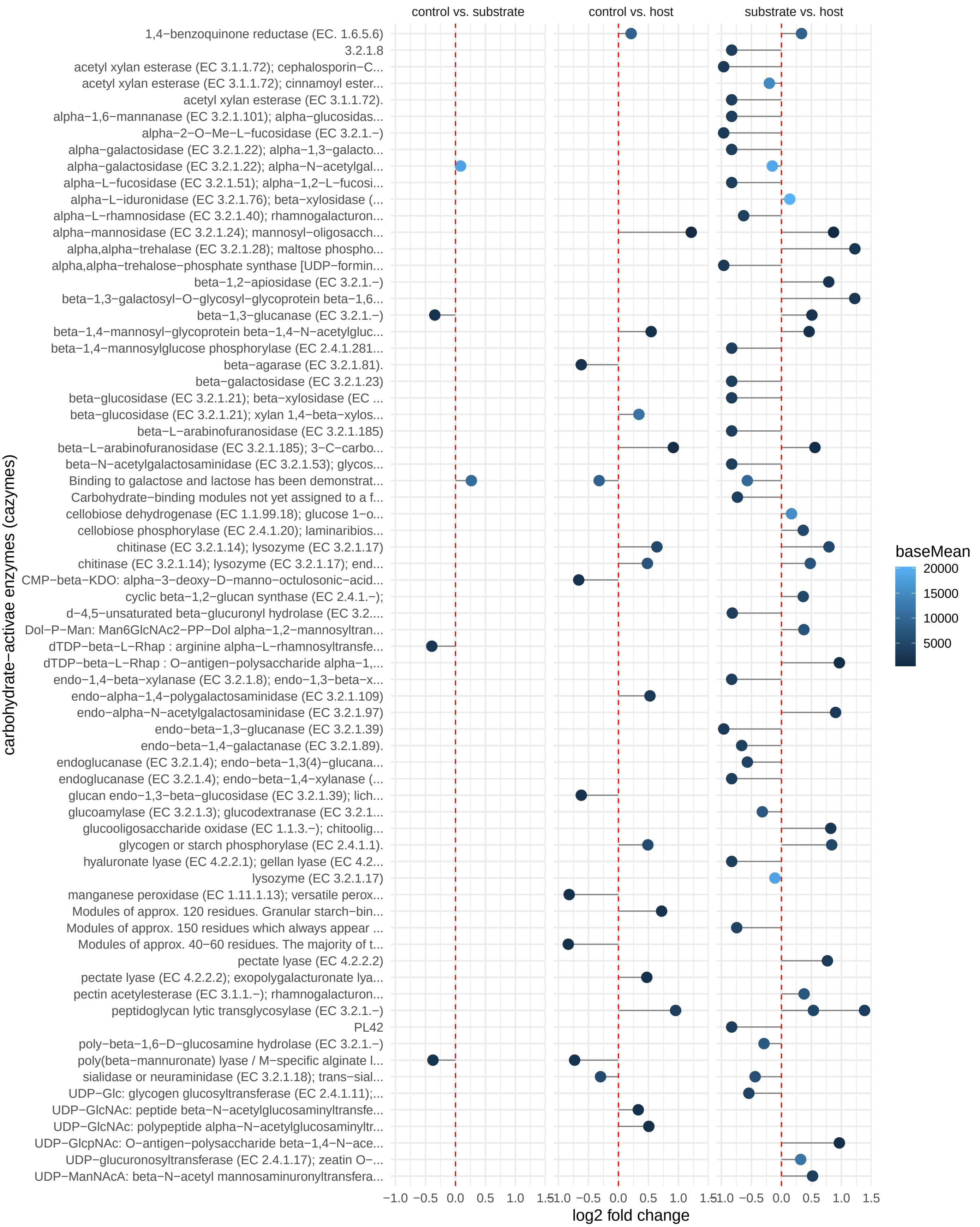

Figure S14: Enriched carbohydrate-active enzymes. Differential abundance analysis was used to compare the abundances of between samples sources.

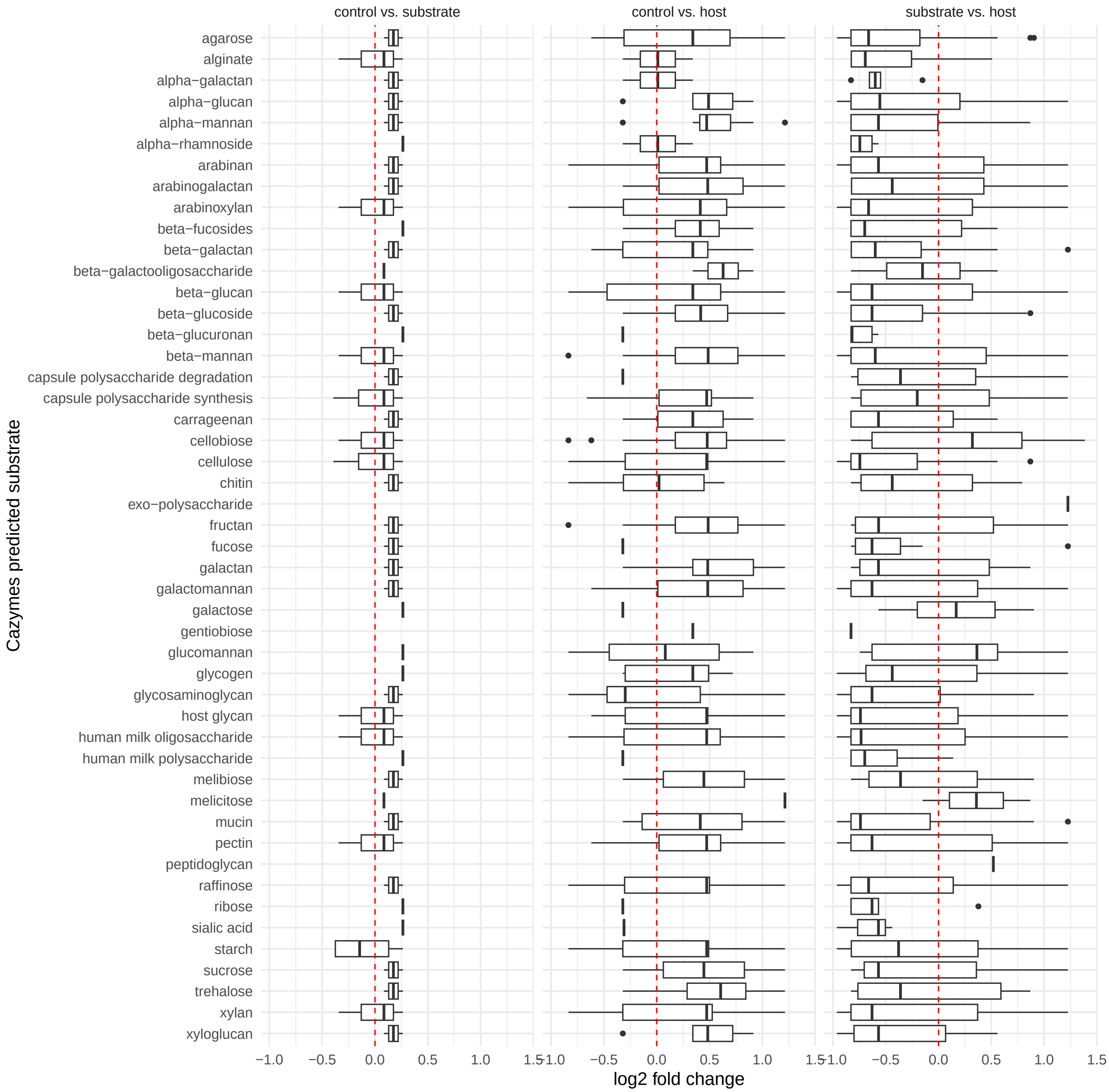

Figure S15: Enriched carbohydrate-active enzymes. Differential abundance analysis was used to compare the abundances of between samples sources.

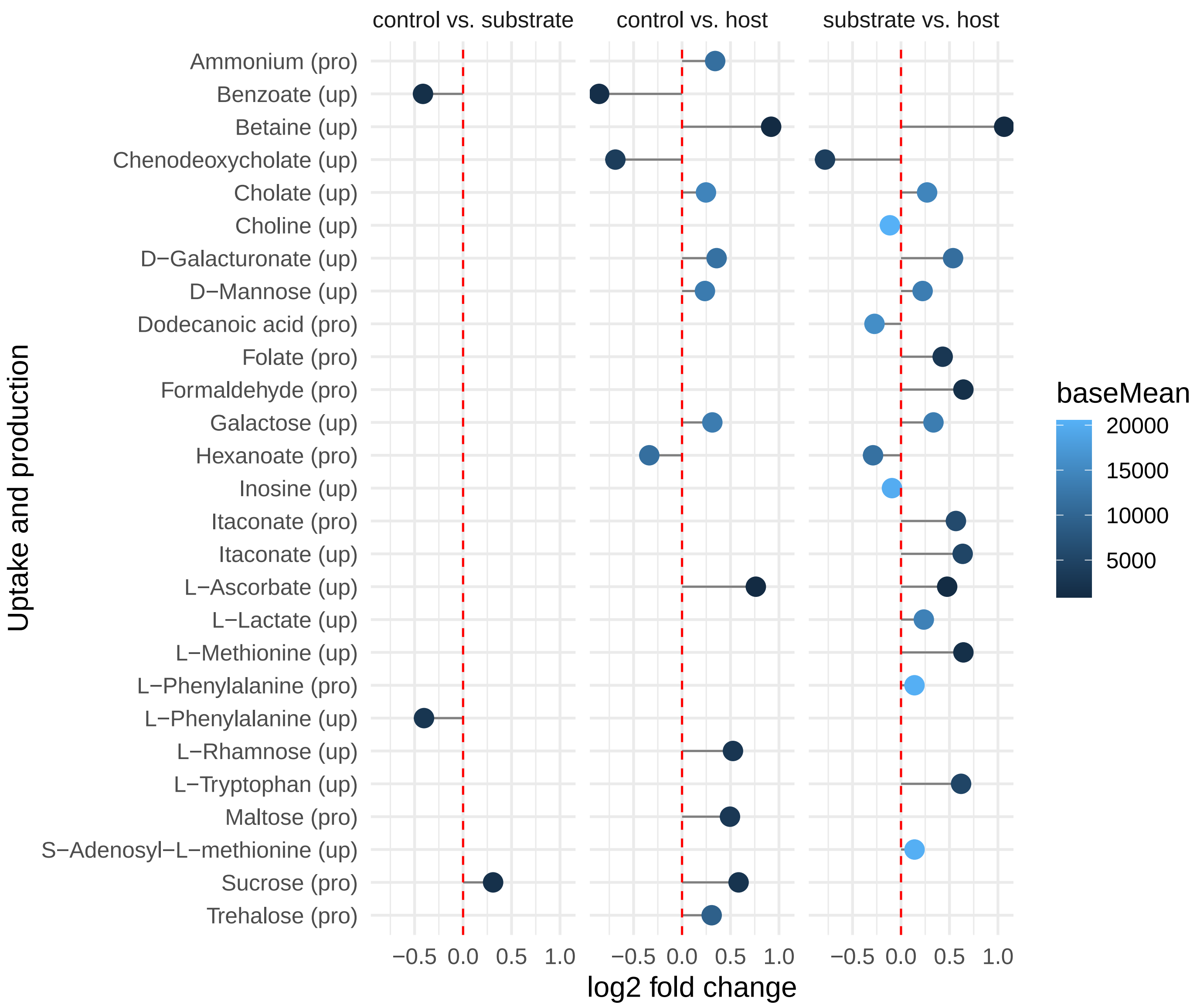

Figure S16: Enriched carbon/electron source (up) and metabolic by-products (pro) predicted from metabolic models using gapseq. Differential abundance analysis was used to compare the abundances of between samples sources.

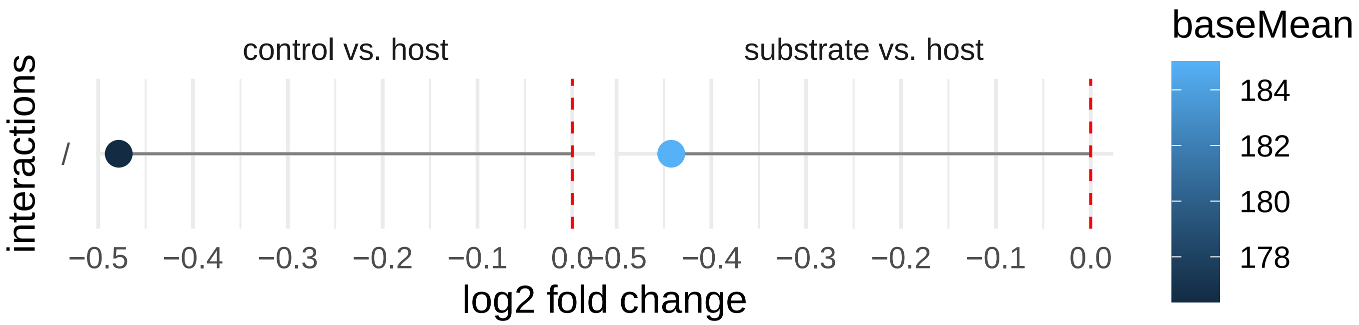

Figure S17: Enriched pair-wise interactions inferred from metabolic models by single vs. co-growth simulations and growth rate comparison. "/" indicates neutral interactions. Differential abundance analysis was used to compare the abundance of between samples sources.

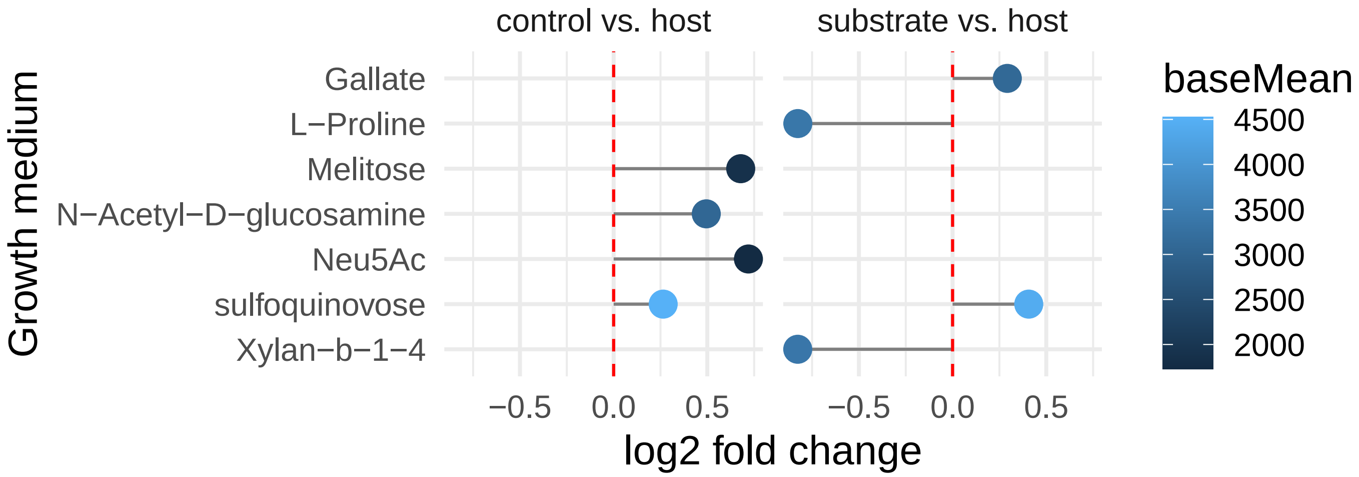

Figure S18: Enriched growth medium compounds predicted from metabolic models by gapseq. Abbreviation: N-acetylneuraminate (Neu5Ac). Differential abundance analysis was used to compare the abundance of between samples sources.

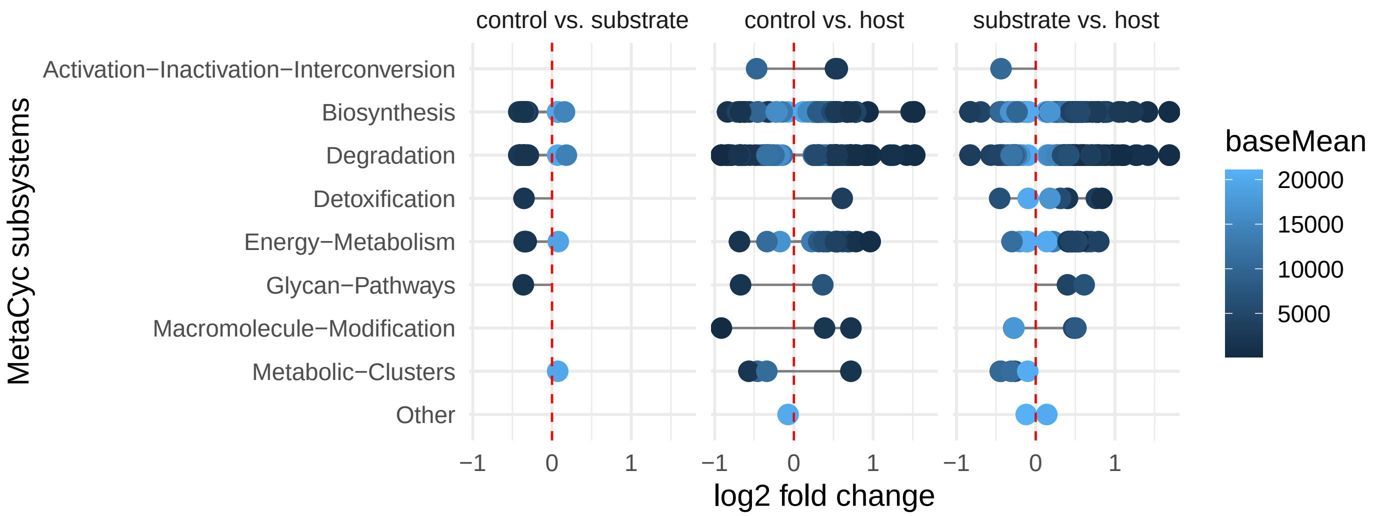

Figure S19: Enriched subsystem for MetaCyc for pathways predicted by gapseq. Differential abundance analysis was used to compare the abundance of between samples sources.

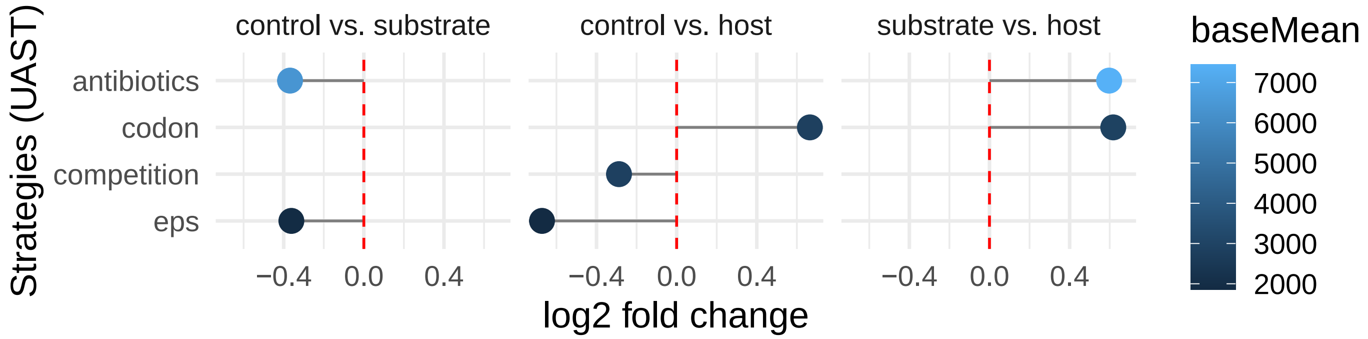

Figure S20: Enriched adaptive strategies and related traits as defined in the universal adaptive strategies (UAST) framework and as applied in a former study [37]. Differential abundance analysis was used to compare the abundance of between samples sources.

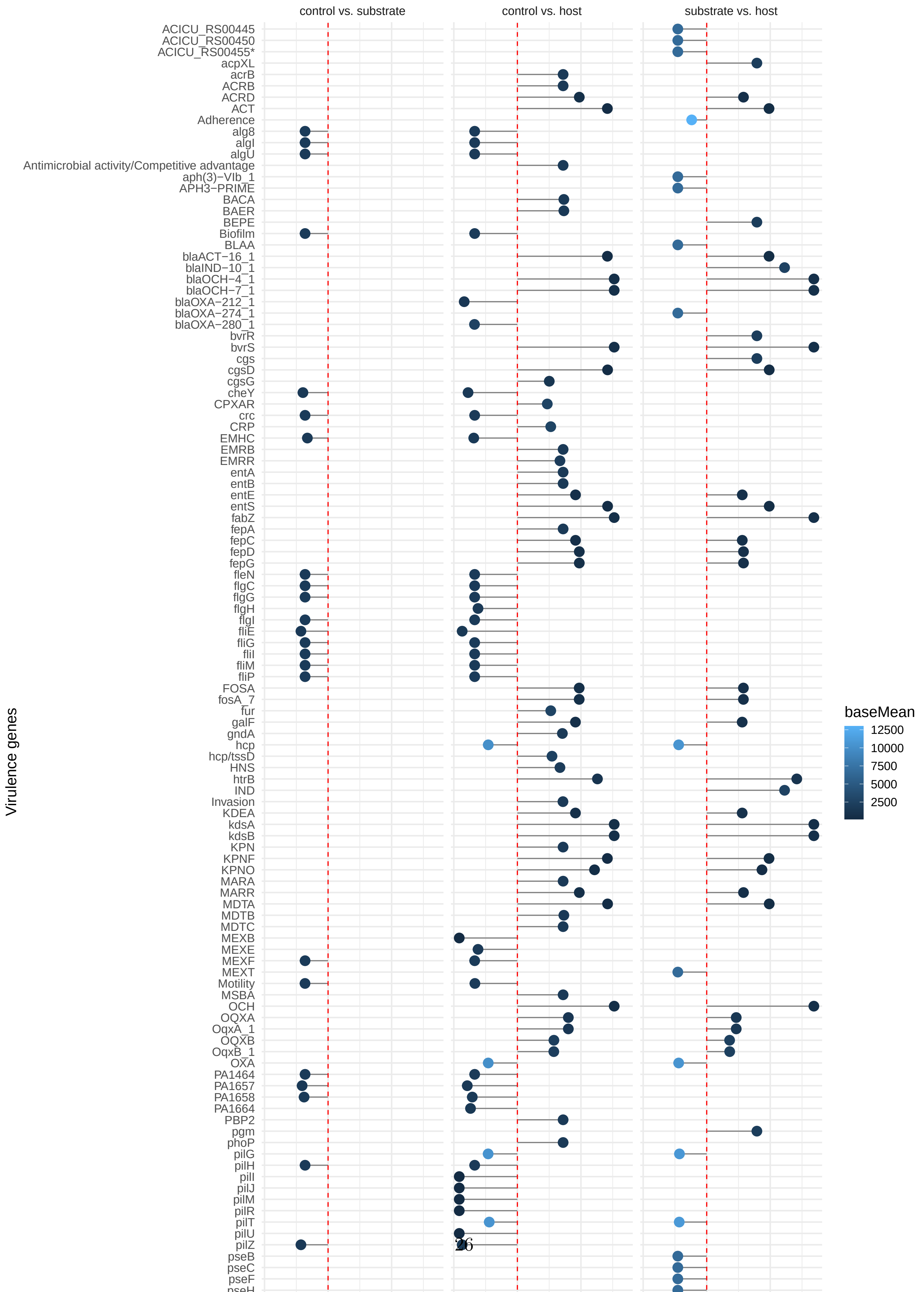
control vs. substrate
control vs. host
substrate vs. host
Virulence genes
ACICU_RS00445
ACICU_RS00450
ACICU_RS00455*
acpXL
acrB
ACRB
ACRD
ACT
Adherence
alg8
algI
algU
Antimicrobial activity/Competitive advantage
aph(3)−VIb_1
APH3−PRIME
BACA
BAER
BEPE
Biofilm
BLAA
blaACT−16_1
blaIND−10_1
blaOCH−4_1
blaOCH−7_1
blaOXA−212_1
blaOXA−274_1
blaOXA−280_1
bvrR
bvrS
cgs
cgsD
cgsG
cheY
CPXAR
crc
CRP
EMHC
EMRB
EMRR
entA
entB
entE
entS
fabZ
fepA
fepC
fepD
fepG
fleN
flgC
flgG
flgH
flgI
fliE
fliG
fliI
fliM
fliP
FOSA
fosA_7
fur
galF
gndA
hcp
hcp/tssD
HNS
htrB
IND
Invasion
KDEA
kdsA
kdsB
KPN
KPNF
KPNO
MARA
MARR
MDTA
MDTB
MDTC
MEXB
MEXE
MEXF
MEXT
Motility
MSBA
OCH
OQXA
OqxA_1
OQXB
OqxB_1
OXA
PA1464
PA1657
PA1658
PA1664
PBP2
pgm
phoP
pilG
pilH
pilI
pilJ
pilM
pilR
pilT
pilU
pilZ
pseB
pseC
pseF
pseH
baseMean
12500
10000
7500
5000
2500

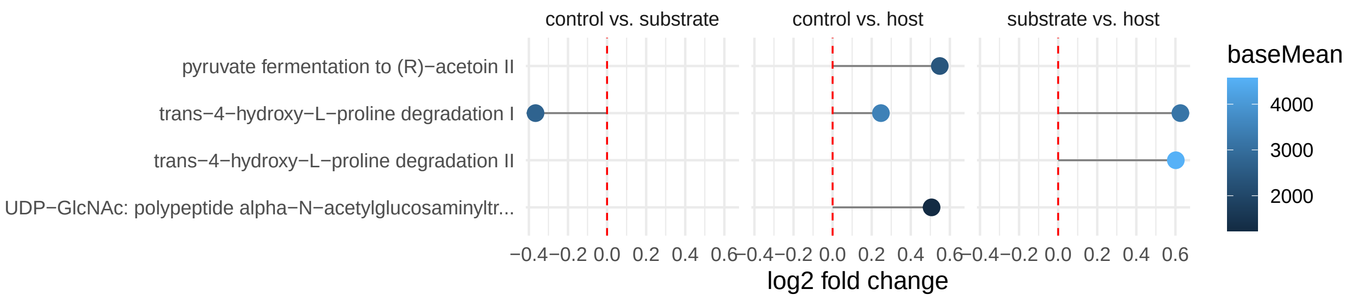

Figure S22: Enriched MetaCyc pathways for hydroxy-proline degradation and acetoin fermentation that we identified previously to be associated with worm fitness and bacterial load [37]. Differential abundance analysis was used to compare the abundance of between samples sources.

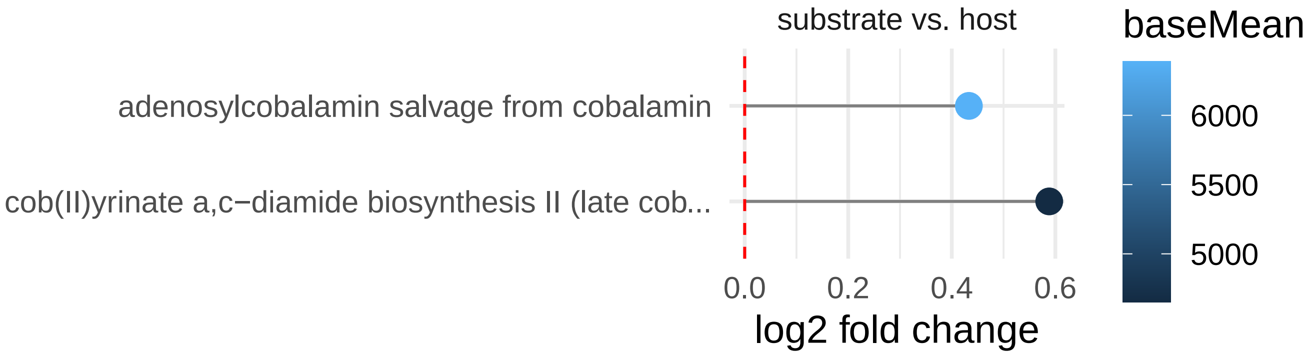

Figure S23: Enriched MetaCyc pathways involving vitamin B12. Differential abundance analysis was used to compare the abundance of between samples sources.

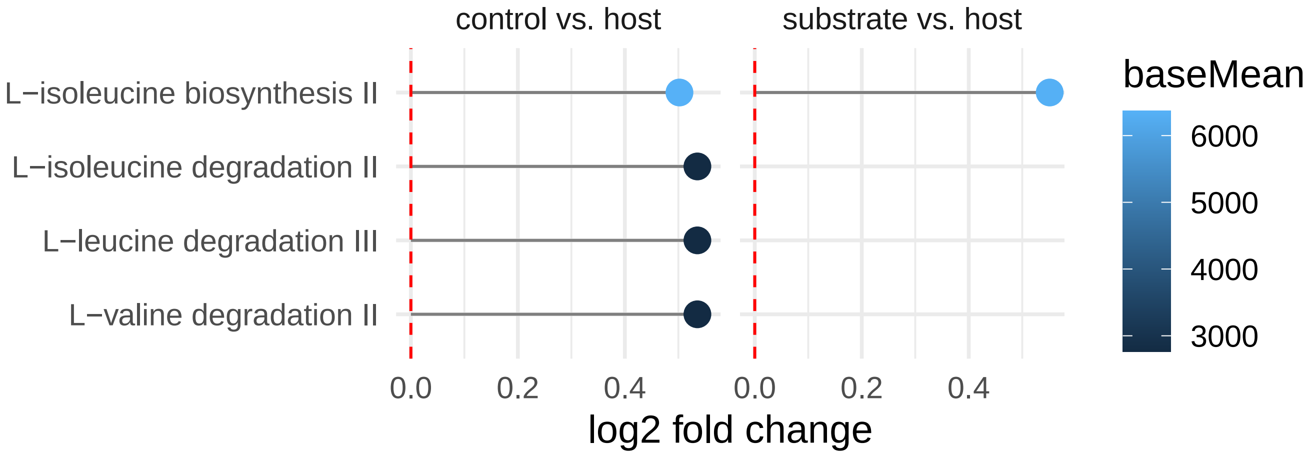

Figure S24: Enriched MetaCyc pathways relevant for branched-chain amino acids metabolism. Differential abundance analysis was used to compare the abundance of between samples sources.

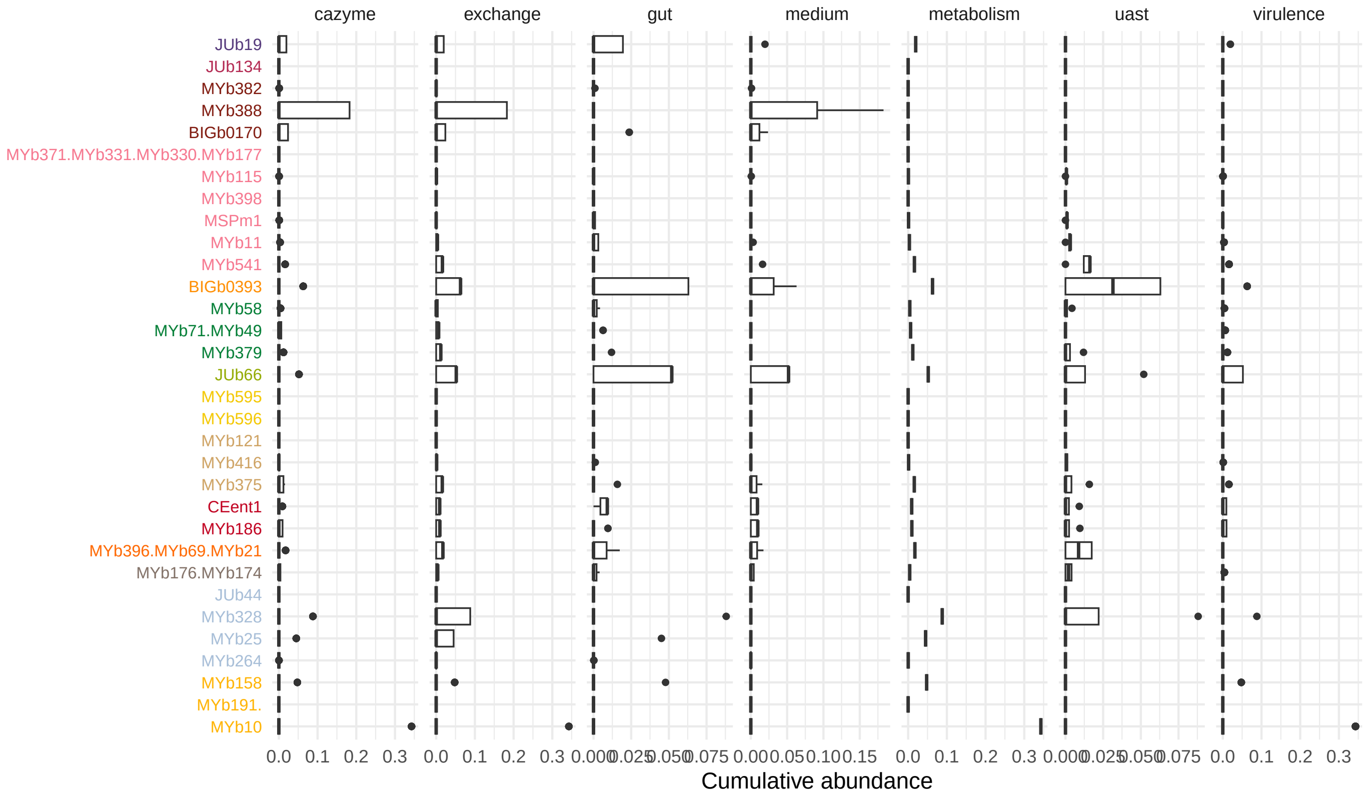

Figure S25: Contribution of microbial species to predicted functions. The attribution of individual taxa to predicted functions was determined by summing up the relative abundances for all species predicted to have the respective function.

### 11 Software

| Program | Version | Parameter |
|---|---|---|
| abricate | 0.8 | -db vfdb |
| dbcan | 3.0.2 | prok -c cluster |
| gutSMASH | 1.0.0-1555cd7 | -genefinding-tool prodigal -cb-knownclusters -cb-general -enable-genefunctions |
| R | 4.1 | |
| tNST (R package NST) | 3.1.10 | rand=1000 |
| nst.boot (R package NST) | 3.1.10 | rand=1000 |
| BLAST | 2.10.1+ | perc_identity 99 -qcov_hsp_perc 95 |
| BLAST (blobtool) | 2.10.1+ | -max_target_seqs 1 -max_hsps 1 -evalue 1e-25 |
| PRINSEQ-lite | 0.20.4 | -min_len 20 -ns_max_n 8 -min_qual_mean 15 -trim_qual_left 12 -trim_qual_right 12 |
| MaSuRCA | 3.4.2 | USE_LINKING_MATES = 1, LIMIT_JUMP_COVERAGE = 60, A_PARAMETERS = cgwErrorRate = 0.25, JF_SIZE = 150000000 |
| gutSMASH | 1.0.0-1555cd7 | -genefinding-tool prodigal -cb-knownclusters -enable-genefunctions |
| BacArena | 1.8.2 | Bac: setAllExInf=TRUE, cellweight_sd=0, limit_growth=F |
| | | Arena: time=3, sec_obj="mtf" |
| fastq_illumina_filter | 0.1 | -keep N -vv |
| SPAdes | v3.14.1 | -careful -only-assembler |

Table S12: Programs including the version numbers and the used parameter are listed. Detailed background on the specific applications of the programs can be found in the Materials and Methods section.

#### References

[1] Anton Bankevich et al. "SPAdes: a new genome assembly algorithm and its applications to single-cell sequencing." eng. In: *J Comput Biol* 19.5 (May 2012), pp. 455–477. URL: http://dx.doi.org/10.1089/cmb.2012.0021.

[2] Eugen Bauer et al. "BacArena: Individual-based metabolic modeling of heterogeneous microbes in complex communities". In: *PLOS Computational Biology* 13.5 (2017), pp. 1–22. URL: https://doi.org/10.1371/journal.pcbi.1005544.

[3] Brian Bushnell. *BBMap short read aligner, and other bioinformatic tools*. sourceforge. 2020. URL: https://sourceforge.net/projects/bbmap/.

[4] Benjamin J Callahan et al. "DADA2: High-resolution sample inference from Illumina amplicon data". en. In: *Nat. Methods* 13.7 (July 2016), pp. 581–583.

[5] Christiam Camacho et al. "BLAST+: architecture and applications." In: *BMC Bioinformatics* 10 (2009), p. 421. URL: https://doi.org/10.1186/1471-2105-10-421.

[6] Alessandra Carattoli et al. "In silico detection and typing of plasmids using PlasmidFinder and plasmid multilocus sequence typing". en. In: *Antimicrob. Agents Chemother.* 58.7 (July 2014), pp. 3895–3903.

[7] Lihong Chen et al. "VFDB 2016: hierarchical and refined dataset for big data analysis—10 years on". In: *Nucleic acids research* 44.D1 (2015), pp. D694–D697. URL: https://doi.org/10.1093/nar/gkv1239.

[8] Philipp Dirksen et al. "CeMbio - The Caenorhabditis elegans Microbiome Resource". In: *G3: Genes, Genomes, Genetics* (July 2020), g3.401309.2020. URL: https://doi.org/10.1534/g3.120.401309.

[9] Philipp Dirksen et al. "The native microbiome of the nematode Caenorhabditis elegans: gateway to a new host-microbiome model". In: *BMC Biology* 14.1 (May 2016).

[10] Enrique Doster et al. "MEGARes 2.0: a database for classification of antimicrobial drug, biocide and metal resistance determinants in metagenomic sequence data". en. In: *Nucleic Acids Res.* 48.D1 (Jan. 2020), pp. D561–D569.

[11] Alexander P. Douglass et al. "Coverage-Versus-Length Plots, a Simple Quality Control Step for de Novo Yeast Genome Sequence Assemblies". In: *G3: Genes, Genomes, Genetics* 9.3 (2019), pp. 879–887. eprint: https://www.g3journal.org/content/9/3/879.full.pdf. URL: https://www.g3journal.org/content/9/3/879.

[12] Zachary S L Foster, Thomas J Sharpton, and Niklaus J Grünwald. "Metacoder: An R package for visualization and manipulation of community taxonomic diversity data". en. In: *PLoS Comput. Biol.* 13.2 (Feb. 2017), e1005404.

[13] Bastian V H Hornung, Romy D Zwittink, and Ed J Kuijper. "Issues and current standards of controls in microbiome research". en. In: *FEMS Microbiol. Ecol.* 95.5 (May 2019).

[14] Julia Johnke, Philipp Dirksen, and Hinrich Schulenburg. "Community assembly of the native C. elegans microbiome is influenced by time, substrate and individual bacterial taxa." In: *Environ Microbiol* 22 (2020), pp. 1265–1279. URL: https://doi.org/10.1111/1462-2920.14932.

[15] DR Laetsch and ML Blaxter. "BlobTools: Interrogation of genome assemblies". In: *F1000Research* 6.1287 (2017).

[16] Heng Li. "Aligning sequence reads, clone sequences and assembly contigs with BWA-MEM". In: (2013).

[17] Heng Li et al. "The sequence alignment/map format and SAMtools". In: *Bioinformatics* 25.16 (2009), pp. 2078–2079.

[18] Michael I. Love, Wolfgang Huber, and Simon Anders. “Moderated estimation of fold change and dispersion for RNA-seq data with DESeq2”. In: *Genome Biology* 15.12 (Dec. 2014), p. 550. URL: https://doi.org/10.1186/s13059-014-0550-8.

[19] Paul J McMurdie and Susan Holmes. “phyloseq: an R package for reproducible interactive analysis and graphics of microbiome census data”. en. In: *PLoS One* 8.4 (Apr. 2013), e61217.

[20] Alla Mikheenko et al. “Versatile genome assembly evaluation with QUAST-LG.” In: *Bioinformatics* 34 (2018), pp. i142–i150. URL: https://doi.org/10.1093/bioinformatics/bty266.

[21] Daliang Ning et al. “A general framework for quantitatively assessing ecological stochasticity”. In: *Proceedings of the National Academy of Sciences* 116.34 (Aug. 2019), pp. 16892–16898. URL: https://doi.org/10.1073/pnas.1904623116.

[22] Andrei Papkou et al. “The genomic basis of Red Queen dynamics during rapid reciprocal host–pathogen coevolution”. In: *Proceedings of the National Academy of Sciences* 116.3 (2019), pp. 923–928.

[23] Victòria Pascal Andreu et al. “The gutSMASH web server: automated identification of primary metabolic gene clusters from the gut microbiota”. In: *Nucleic Acids Research* 49.W1 (May 2021), W263–W270. eprint: https://academic.oup.com/nar/article-pdf/49/W1/W263/38842391/gkab353.pdf. URL: https://doi.org/10.1093/nar/gkab353.

[24] Robert Schmieder and Robert Edwards. “Quality control and preprocessing of metagenomic datasets”. In: *Bioinformatics* 27.6 (Jan. 2011), pp. 863–864. eprint: https://academic.oup.com/bioinformatics/article-pdf/27/6/863/646767/btr026.pdf. URL: https://doi.org/10.1093/bioinformatics/btr026.

[25] Thorsten Seemann. *ABRicate: Mass screening of contigs for antimicrobial and virulence genes*. github. 2020. URL: https://github.com/tseemann/abricate.

[26] Torsten Seemann. *Shovill: Assemble bacterial isolate genomes from Illumina paired-end reads*. github. 2020. URL: https://github.com/tseemann/shovill.

[27] Michael Sieber et al. “Neutrality in the Metaorganism.” In: *PLoS Biol.* 17 (2019), e3000298. URL: https://doi.org/10.1371/journal.pbio.3000298.

[28] William T Sloan et al. “Quantifying the roles of immigration and chance in shaping prokaryote community structure”. In: *Environmental Microbiology* 8.4 (2006), pp. 732–740.

[29] Alexandre Souvorov, Richa Agarwala, and David J Lipman. “SKESA: strategic k-mer extension for scrupulous assemblies”. en. In: *Genome Biol.* 19.1 (Oct. 2018), p. 153.

[30] Theresa Stiernagle. “Maintenance of C. elegans”. In: *WormBook* (2006).

[31] Daniel Straub et al. “Interpretations of environmental microbial community studies are biased by the selected 16S rRNA (gene) amplicon sequencing pipeline”. en. In: *Front. Microbiol.* 11 (Oct. 2020), p. 550420.

[32] Ole Tange. *Gnu Parallel 2018*. en. Zenodo, 2018. URL: https://zenodo.org/record/1146014.

[33] Ryan R Wick et al. “Unicycler: Resolving bacterial genome assemblies from short and long sequencing reads.” In: *PLoS Comput. Biol.* 13 (2017), e1005595. URL: https://doi.org/10.1371/journal.pcbi.1005595.

[34] Ea Zankari et al. “Identification of acquired antimicrobial resistance genes”. en. In: *J. Antimicrob. Chemother.* 67.11 (Nov. 2012), pp. 2640–2644.

[35] Han Zhang et al. “dbCAN2: a meta server for automated carbohydrate-active enzyme annotation”. In: *Nucleic Acids Research* 46.W1 (2018), W95–W101. URL: http://dx.doi.org/10.1093/nar/gky418.

[36] A. V. Zimin et al. “The MaSuRCA genome assembler”. In: *Bioinformatics* 29.21 (Nov. 2013), pp. 2669–2677. URL: http://dx.doi.org/10.1093/bioinformatics/btt476.

[37] Johannes Zimmermann* et al. "The functional repertoire contained within the native microbiota of the model nematode *Caenorhabditis elegans*." In: *ISME Journal* 14.1 (2020), pp. 26–38. URL: https://doi.org/10.1038/s41396-019-0504-y.
